# vmTracking: Virtual Markers Overcome Occlusion and Crowding in Multi-Animal Pose Tracking

**DOI:** 10.1101/2024.02.07.579241

**Authors:** Hirotsugu Azechi, Susumu Takahashi

**Affiliations:** Laboratory of Cognitive and Behavioral Neuroscience, Graduate School of Brain Science, Doshisha University, Kyotanabe, 610-0394, Japan

**Keywords:** multi-animal tracking, virtual marker, occlusion, crowding, DeepLabCut, SLEAP

## Abstract

In multi-animal tracking, addressing occlusion and crowding is crucial for accurate behavioral analysis. However, in situations where occlusion and crowding generate complex interactions, achieving accurate pose tracking remains challenging. Therefore, we introduced Virtual marker tracking (vmTracking), which uses virtual markers for individual identification. Virtual markers are labels derived from conventional markerless multi-animal tracking tools, such as multi-animal DeepLabCut (maDLC) and Social LEAP Estimate Animal Poses (SLEAP). Unlike physical markers, virtual markers exist only within the video and attribute features to individuals, enabling consistent identification throughout the entire video while keeping the animals markerless in reality. Using these markers as cues, annotations were applied to multi-animal videos, and tracking was conducted with single-animal DeepLabCut (saDLC) and SLEAP’s single-animal method. vmTracking minimized manual corrections and annotation frames needed for training, efficiently tackling occlusion and crowding. Experiments tracking multiple mice, fish, and human dancers confirmed vmTracking’s variability and applicability. These findings could enhance the precision and reliability of tracking methods used in the analysis of complex naturalistic and social behaviors in animals, providing a simpler yet more effective solution.

## Introduction

The study of animal behavior is crucial to advance our understanding of biological, physiological, and neurological processes. Precise simultaneous pose tracking of multiple animals is essential for analyzing naturalistic and social behaviors across species. Recent advancements in machine learning have produced high-precision and versatile single-animal pose tracking tools, such as DeepLabCut (DLC) [1] and LEAP Estimates Animal Poses (LEAP) [2], allowing detailed analysis of complex postures and movements across a wide range of species [3–6]. Building on this progress, multi-animal tracking tools such as multi-animal DeepLabCut (maDLC) [7] and Social LEAP (SLEAP) [8] have been introduced. These tools use methods such as Kalman filters [9] or optical flow [10] to predict movements and facilitate frame-to-frame identification. However, despite their effectiveness in simpler scenarios, these tools face substantial challenges in multi-animal contexts, particularly with issues such as occlusion, where animals obscure each other, and crowding, where individuals are densely clustered. Despite ongoing research and development efforts to resolve issues of occlusion and crowding [11,12] these problems remain unsolved, hindering accurate multi-animal pose tracking. Currently, research using these tools has succeeded in tracking simple postures such as head direction [13–15], but it is not advanced enough to accurately track detailed postures over extended periods.

While automated systems face challenges in achieving high accuracy in animal tracking, human involvement in reviewing, correcting, and providing feedback on the results can enhance accuracy. Thus, human-in-the-loop machine learning, which integrates human judgment into the machine-learning process, can enhance animal-tracking methods [16]. Currently, in simple scenarios without occlusion or crowding, accurate multi-animal pose tracking can be achieved with minimal human intervention. However, in situations where occlusion and crowding generate complexinteractions, the need for human intervention increases significantly. Achieving accurate pose tracking still remains challenging.

In this study, to address the challenges of occlusion and crowding, we incorporated human intervention during the tracking process to introduce virtual markers into videos of markerless subjects. These markers provide individual identification features without relying on physical markers, which can potentially have unforeseen effects on animal behavior and may inhibit natural behaviors. Through tracking experiments with mice, fish, and humans, we demonstrate that our vmTracking approach with virtual markers enhances the accuracy of machine learning predictions in occlusion and crowding while requiring minimal manual human intervention. The vmTracking provides a robust framework for observing and analyzing social interactions among animals in their natural environments, potentially expanding the boundaries of behavioral science and offering deep insights into collective animal behavior.

## Results

The concept of vmTracking is based on the observation that multiple individuals can be tracked with single-animal DeepLabCut (saDLC) if they are visually distinguishable (e.g., black and white mice) [17]. By extending this idea, it may be possible to track visually indistinguishable markerless animals by assigning certain identifiers as a marker. For this purpose, we decided to use the labels obtained from conventional markerless multi-animal tracking as “virtual” markers, which facilitated tracking of multiple animals while keeping them markerless in reality (Fig 1). Thus, in vmTracking, two separate tracking projects were conducted for different purposes: assigning virtual markers and tracking those markers.

**Fig 1.**
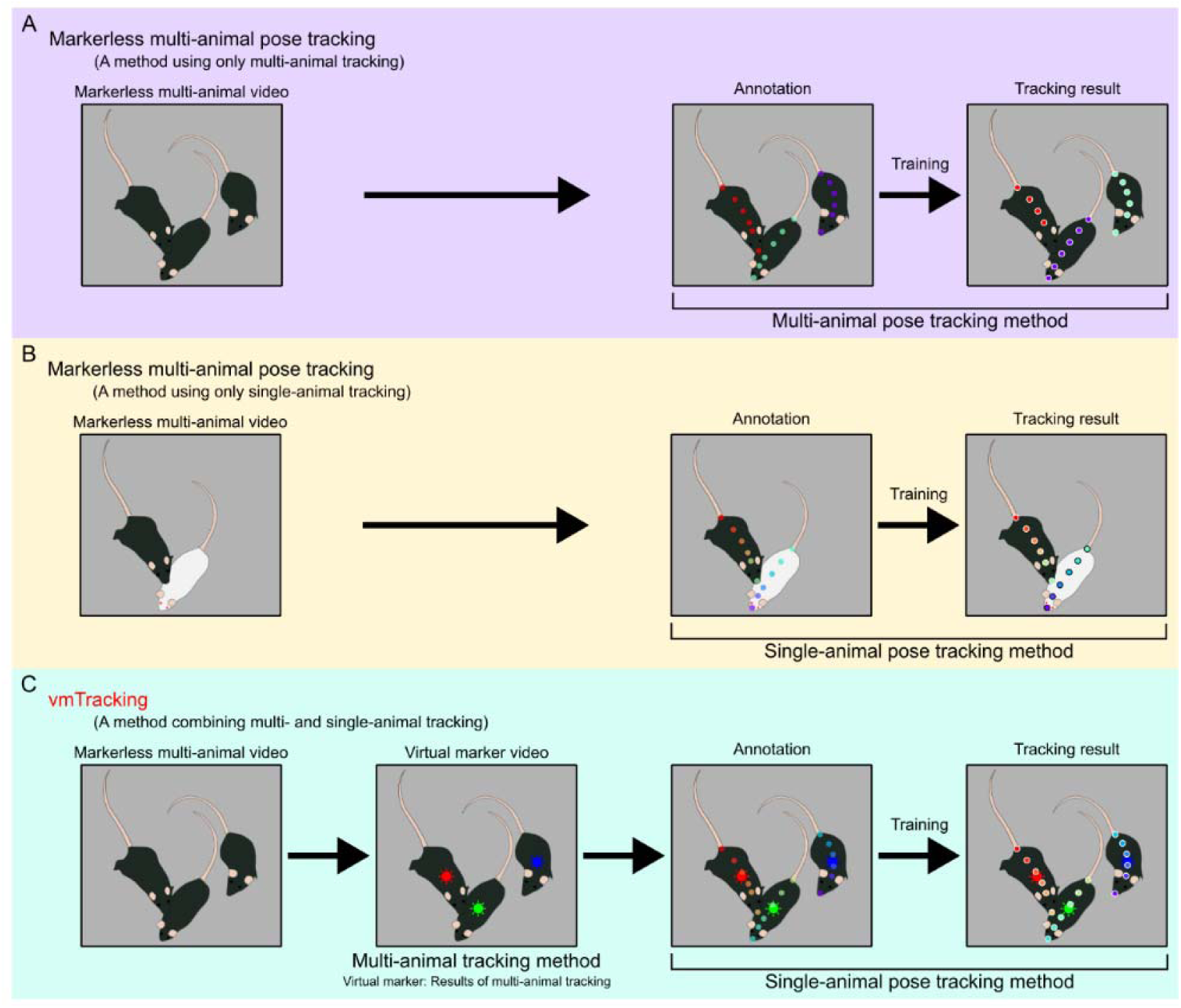
vmTracking concept. (A) In traditional markerless multi-animal pose tracking, the poses of multiple animals that look the same can be tracked. However, ID switches may occur during occlusion or crowding events, and predicted keypoints can be missing even in non-occluded regions. (B) Even for markerless animals, when tracking the poses of two mice with different colors, a single-animal tracking method can track both mice by using the color difference as a feature. In particular, with DeepLabCut, missing predicted keypoints, which are sometimes observed in multi-animal pose tracking methods, do not occur. However, this method is only applicable under limited conditions, such as when the animals have different colors. (C) vmTracking integrates these features. First, a video visualizing the results obtained from markerless multi-animal tracking is created. Then, by applying a single-animal pose tracking method to the labels in the video as virtual identification markers, it performs pose tracking of multiple animals that are physically markerless.

First, markerless multi-animal pose tracking was performed, and then a labeled video was created by outputting the tracking results. These labels, which were only observed within the video, were termed “virtual markers” and were used for consistent individual identification. Videos of multiple animals marked with these virtual markers were then analyzed using a single-animal pose tracking tool (either saDLC or SLEAP’s single-animal method, LEAP). During annotation, each animal was labeled with a consistent ID across frames based on the virtual marker. The accuracy of virtual marker tracking with either saDLC (vmT-DLC) or LEAP (vmT-LEAP) were compared to those of traditional markerless multi-animal pose tracking tools (maDLC or SLEAP) for metrics based on ground truth (S1 Fig).

### Tracking black mice against black backgrounds

To evaluate the effectiveness of vmTracking in low-contrast environments, where individual identification is particularly challenging due to occlusion and crowding, an experiment was conducted with 5 C57BL/6J mice on a black background, and 12 scenes with occlusion and crowding were evaluated. A brief outline of the experiment is shown in Fig 2A. Each mouse was tracked at six keypoints. Markerless multi-animal pose tracking was performed using maDLC or SLEAP (Fig 2B and 2C). Based on intermediate data from refining the markerless tracking results, we created a video visualizing two keypoints as virtual markers using SLEAP (Fig 2B and 2D) and conducted vmT-DLC and vmT-LEAP (Fig 2E and S1 and S2 Video). The final total number of annotated frames for maDLC and vmT-DLC was 2,097 and 1,196, respectively. The results for SLEAP and vmT-LEAP were based on training and predictions using annotation data imported from maDLC and vmT-DLC.

**Fig 2.**
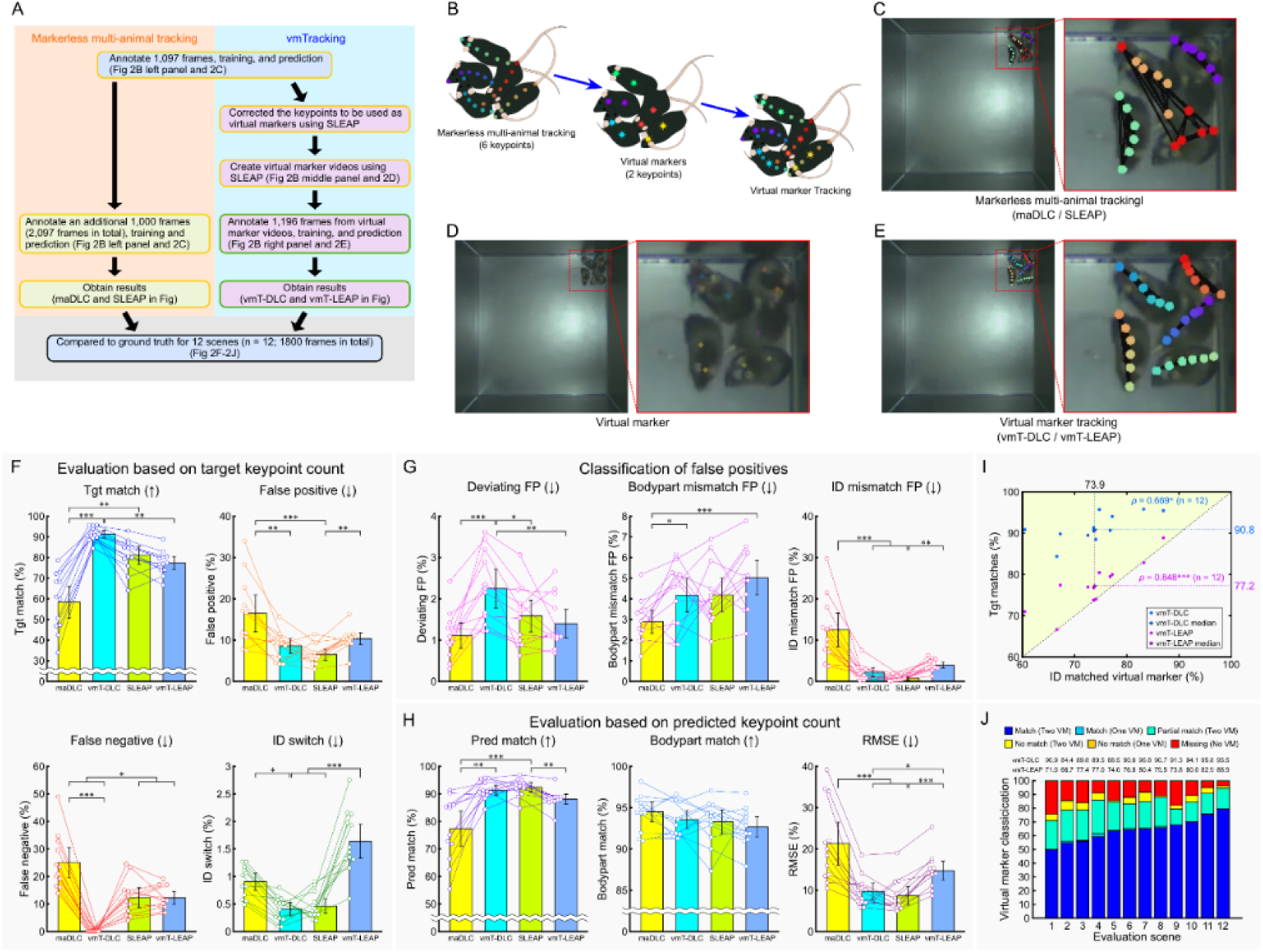
Comparison of markerless multi-animal pose tracking and vmTracking for tracking five mice. (A) Overview of the validation process. The yellow box in the schematic represents the processes using multi-animal tracking tools, while the green box represents the processes using single-animal tracking tools. (B) Schematic of virtual marker creation. Six keypoints per mouse were tracked; two designated as virtual markers. (C–E) Snapshots of markerless multi-animal pose tracking in occlusion-crowded scenes (C), virtual markers (D), and virtual marker tracking (E). Red lines on the right panel represent the enlarged view of the red-lined box in the left panel. (F–H) Various percentage-based metrics were compared across multi-animal DeepLabCut (maDLC), virtual marker tracking with DeepLabCut (vmT-DLC), social LEAP (SLEAP), and virtual marker tracking with SLEAP (vmT-LEAP) based on manually generated ground truth (GT) data. (F) Evaluation was conducted as a percentage of the target keypoints, with the total number of GT keypoints as the denominator. The percentages of matches (Tgt match), false negatives, false positives, and ID switches were calculated based on the number of GT keypoints. (G) False positives from (F) were classified into deviations exceeding the threshold from all GT (Deviating FP), mismatches in predicted body parts (Bodypart mismatch FP), and cases where the predicted body part was correct but the ID was incorrect (ID mismatch FP). (H) Evaluation was conducted as a percentage of the predicted keypoints, excluding FNs, using the total predicted keypoints as the denominator, and the metrics included matches (Pred match), body part matches (Bodypart match), irrespective of ID, and root mean square error (RMSE). Arrows indicate whether higher or lower values are better. Data are presented as mean ± 95% confidence interval (n = 12 scenes). Statistical analysis was performed using the Friedman test with Bonferroni correction. (I) Scatter plots illustrating the relationship between ID-matched virtual markers and Tgt matches for vmT-DLC and vmT-LEAP. Plots in the upper-left (light yellow-green) area indicate cases where Tgt matches exceeded the ID-matched virtual markers. Filled circles represent individual scenes, and the stars signify their median. Correlation analysis was conducted using Spearman’s rank correlation (n = 12 scenes). All statistical significance levels were set at p < 0.05, *: p < 0.05, **: p < 0.01, ***: p < 0.001. (J) Stacked bar graph showing the proportion of virtual marker assignment patterns in each scene. Virtual markers were classified as follows: when two virtual markers were assigned to a single mouse, both IDs matched (Match (Two VM)); one virtual marker was missing and the other’s ID matched (Match (One VM)); one virtual marker’s ID matched while the other’s ID did not match (Partial match (Two VM)); neither ID matched (No match (Two VM)); one virtual marker was missing and the other’s ID did not match (No match (One VM)); and both virtual markers were missing (Missing (Two VM)). The proportions of each category are displayed by scene. The values for vmT-DLC and vmT-LEAP above the figure represent the Tgt match values for each scene.

A comparison of the number of target points (equivalent to the GT count) (Friedman test with Bonferroni correction, n = 12) revealed that vmT-DLC showed improved performance over maDLC in Tgt match (p < 0.001), false positives (FPs) (p < 0.01), false negatives (FNs) (p < 0.001), and ID switches (p < 0.05) (Fig 2F). However, when the FPs were further classified, vmT-DLC showed significantly better performance in ID mismatch FPs (p < 0.001) while maDLC showed significantly better performance in Deviating FPs (p < 0.001) and Bodypart mismatch FPs (p < 0.05) (Fig 2G). A comparison based on predicted keypoints revealed that vmT-DLC achieved significantly higher Pred match (p < 0.01) and lower RMSE (p < 0.001) values than maDLC, while no difference was observed for Bodypart match (Fig 2H). Here, as vmT-DLC had 0% FNs (Fig 2F), its Tgt match and Pred match values were the same. Conversely, the only metric in which vmT-DLC was significantly better than SLEAP was FNs (p < 0.05), while SLEAP was better than vmT-DLC in Deviating FPs (p < 0.05). In contrast to vmT-DLC, the only metric in which vmT-LEAP performed significantly better than maDLC or SLEAP was FNs (p < 0.05) against maDLC (Fig 2F), with SLEAP demonstrating better results than vmT-LEAP in FPs (p < 0.01), ID switches (p < 0.001), ID mismatch FPs (p < 0.01), Pred match (p < 0.01), and RMSE (p < 0.001). No metrics were found in which vmT-LEAP outperformed SLEAP (Fig 2F–2H). Notably, no significant differences were found for Bodypart match, the least stringent matching criterion, across all methods (Fig 2H).

The relationship between virtual marker accuracy (ID matched virtual marker) and Tgt match in vmTracking showed a significant positive correlation for both vmT-DLC and vmT-LEAP (vmT-DLC: *ρ* = 0.669, p < 0.05; vmT-LEAP: *ρ* = 0.848, p < 0.001; Fig 2I). Furthermore, we examined the relationship between virtual marker pattern classification and Tgt match within each scene. Analysis of individual scenes revealed that the Tgt match showed higher values than the proportion of assigned virtual markers with at least one matching ID (sum of Match and Partial match) (Fig 2J), particularly for vmT-DLC.

In summary, under conditions of tracking five black mice against a black background, vmT-DLC outperformed maDLC in terms of matches despite a higher number of certain FPs; however, it did not outperform SLEAP. Conversely, vmT-LEAP did not exceed the performance of either maDLC or SLEAP. Moreover, vmTracking generally appeared to improve the tracking accuracy as the ID match rate increased. Specifically, vmT-DLC suggests that even for mice with non-matching virtual markers (No match) or missing virtual markers (Missing), correct predictions were made for some of those mice.

### Tracking of black mice against white backgrounds

To validate the performance of vmTracking under different conditions, an experiment used three mice against a white background, focusing on scenarios classified as either occlusion-crowded (OC) (Fig 3) or non-occlusion-crowded (nOC) (Fig S3). This distinction facilitated an assessment of vmTracking across different levels of complexity within the same environment. A brief outline of the experiment is shown in Figs 3A and S3A. As in the five-mice tracking experiment, six keypoints were assigned per mouse for markerless multi-animal pose tracking (Fig 3B and 3C, left, and S3B and S3C Fig, left). Subsequently, videos were created to visualize two keypoints as virtual markers (Fig 3B and 3C, middle, and S3B and S3C Fig, middle), and virtual marker tracking was conducted (Fig 3B and 3C, right, and S3B and S3C Fig (right), and S1, S3, and S4 Video). Additionally, to simplify the creation of virtual markers, we attempted maDLC by setting only the two points used as virtual markers as keypoints. However, this approach failed to stitch tracklets together, resulting in the inability to obtain coordinate data; consequently, it was impossible to create virtual marker videos. Consequently, virtual marker tracking was performed using the virtual marker videos created from six-point tracking. The results for maDLC and vmT-DLC were obtained from a final total of 3,886 annotated frames and 1,167 annotated frames, respectively. The results for SLEAP and vmT-LEAP were based on training and predictions using annotation data imported from maDLC and vmT-DLC.

**Fig 3.**
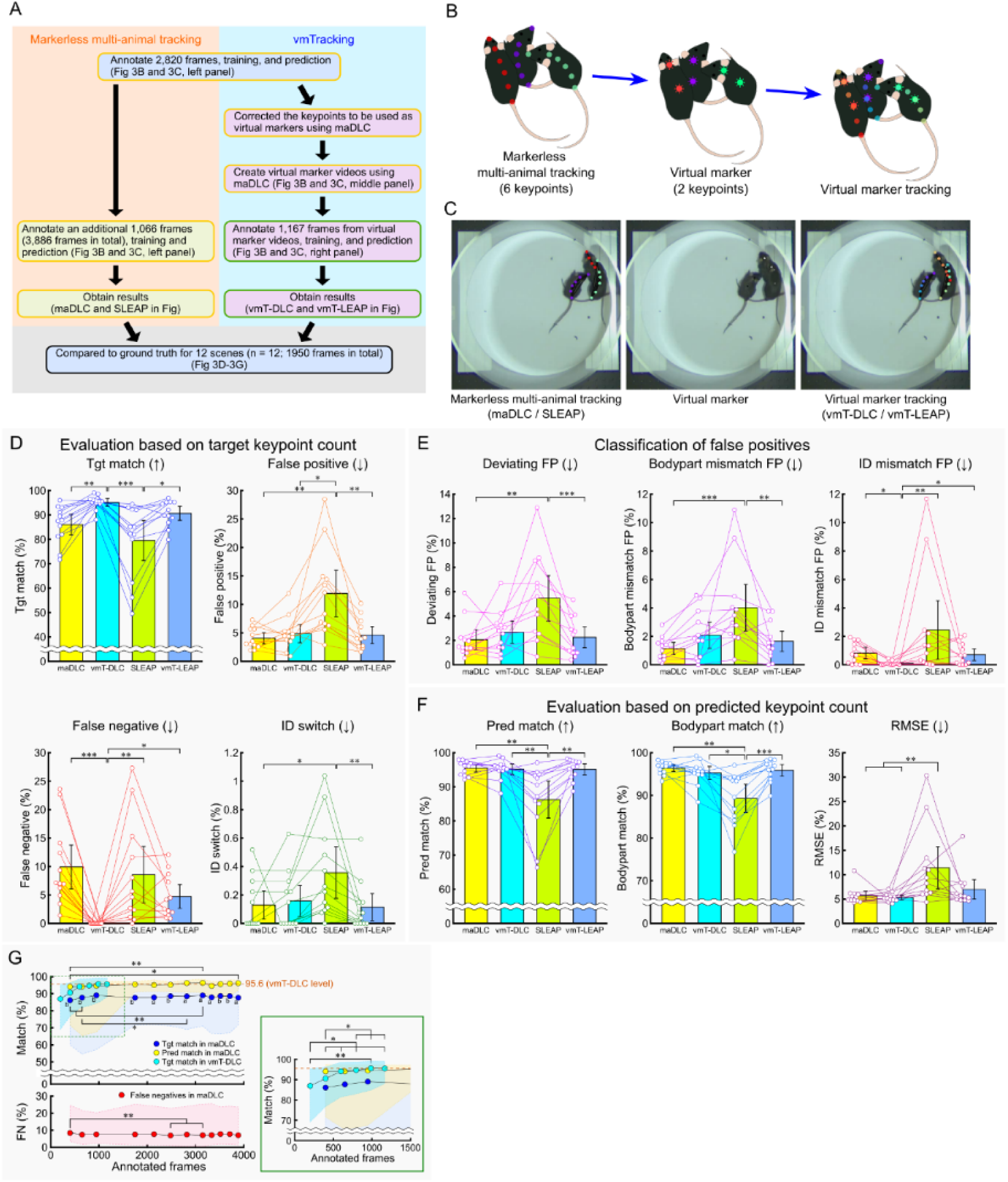
Comparison of markerless multi-animal pose tracking and vmTracking in an occlusion-crowded scene of three mice. (A) Overview of the verification process. The yellow box in the schematic represents the processes using multi-animal tracking tools, while the green box represents the processes using single-animal tracking tools. (B) Schematic of virtual marker creation. Six keypoints per mouse were tracked, with two designated as virtual markers. (C) Examples of markerless multi-animal pose tracking (left), virtual marker (middle), and virtual marker tracking (right) in an occlusion-crowded scene. (D–F) Various percentage-based metrics were compared across multi-animal DeepLabCut (maDLC), virtual marker tracking with DeepLabCut (vmT-DLC), Social LEAP (SLEAP), and virtual marker tracking with SLEAP (vmT-LEAP), based on manually generated ground truth (GT) data. (D) The evaluation was conducted as a percentage of the target keypoints, with the total number of GT keypoints as the denominator. The percentages of matches (Tgt match), false negatives, false positives, and ID switches were calculated based on the number of GT keypoints. (E) The false positives from (D) were classified into deviations exceeding the threshold from all GT (Deviating FP), mismatches in predicted body parts (Bodypart mismatch FP), and cases where the predicted body part was correct but the ID was incorrect (ID mismatch FP). (F) The evaluation was conducted as a percentage of the predicted keypoints, excluding FNs, using the total predicted keypoints as the denominator, and the metrics included matches (Pred match), body part matches (Bodypart match), irrespective of ID, and root mean square error (RMSE). Arrows indicate whether higher or lower values are better. Data are shown as mean ± 95% confidence interval (n = 12 scenes). Statistical analysis was performed using the Friedman test with Bonferroni correction. (G) The relationship between the number of annotated frames and the ratios of Tgt match, Pred match, and false negatives (FNs). The plot shows the median for each number of annotation frames. The shaded area represents the range between the minimum and maximum values. The orange dashed line in the figure represents the median of Tgt match for vmT-DLC in (D). The orange dashed line in the figure represents the Tgt matches of vmT-DLC (corresponding to vmT-DLC Tgt match in Fig 3D). The accuracy across annotation frame counts for each metric was compared using the Friedman test with Bonferroni correction. Additionally, for the “Matches” metric, a paired comparison was conducted using the Wilcoxon signed-rank test against the accuracy of vmT-DLC level, followed by Bonferroni correction. All statistical significance levels were set at p < 0.05. *: p < 0.05, **: p < 0.01, a: p < 0.05 compared to vmT-DLC level, b: p < 0.01 compared to vmT-DLC level.

In the comparison of metrics based on the number of target keypoints for OC scenes (Friedman test with Bonferroni correction, n = 12), vmT-DLC performed significantly better than maDLC in Tgt match (p < 0.01) and FNs (p < 0.001) (Fig 3D). When comparing FPs by category, vmT-DLC showed a significant improvement over maDLC in ID mismatch FPs (p < 0.05), although there were no significant differences in Deviating FPs and Bodypart mismatch FPs (Fig 3E). In the comparison based on the predicted keypoint count, there were no significant differences between vmT-DLC and maDLC across all metrics (Fig 3F). Moreover, as the FNs were 0% for vmT-DLC (Fig 3D), the values for Tgt match and Pred match were identical. On the contrary, under these conditions, vmT-DLC was significantly better than SLEAP in Tgt match (p < 0.001), FPs (p < 0.05), FNs (p < 0.01), ID mismatch FPs (p < 0.01), Pred match (p < 0.01), Bodypart match (p < 0.05), and RMSE (p < 0.01) (Fig 3D–3F). Additionally, vmT-LEAP also performed significantly better than SLEAP in Tgt match (p < 0.05), FPs (p < 0.01), ID switches (p < 0.01), Deviating FPs (p < 0.001), Bodypart mismatch FPs (p < 0.01), Pred match (p < 0.01), and Bodypart match (p < 0.001) (Fig 3D–3F). However, no metrics showed vmT-LEAP to be significantly superior to maDLC (Fig 3D–3F).

In the comparison of metrics based on the number of target keypoints for nOC scenes (Friedman test with Bonferroni correction, n = 8), vmT-DLC performed significantly better than maDLC only in FNs (p < 0.05), with no significant differences observed in other metrics (S3D–S3F Fig). When comparing vmT-DLC with SLEAP, vmT-DLC showed significantly better performance in Tgt match (p < 0.001), FPs (p < 0.001), Deviating FPs (p < 0.05), Bodypart mismatch FPs (p < 0.01), Pred match (p < 0.001), Bodypart match (p < 0.001), and RMSE (p < 0.05) (S3D-S3F Fig). vmT-LEAP also performed significantly better than SLEAP in FPs (p < 0.05), Deviating FPs (p < 0.05), Bodypart mismatch FPs (p < 0.01), Pred match (p < 0.05), Bodypart match (p < 0.05), and RMSE (p < 0.05), but no significant differences were observed across any metrics when comparing vmT-LEAP with maDLC (S3D–S3F Fig).

When evaluating both OC and nOC scenes together, a general positive correlation was observed between ID-matched virtual markers and Tgt match results under these conditions, with significant positive correlations for both vmT-DLC and vmT-LEAP (vmT-DLC: *ρ* = 0.547, p < 0.05; S3G Fig, left, vmT-LEAP: *ρ* = 0.744, p < 0.001; S3G Fig, right). Furthermore, the relationship between the classification of virtual marker patterns and Tgt match within each scene was examined.

Similar to the findings of the five-mice tracking experiment, particularly in the OC scenes of vmT-DLC, the Tgt match was higher than the proportion of virtual markers assigned with at least one ID match (the sum of Match and Partial match) in some scenes (S3H Fig). On the contrary, in the nOC scenes, the proportion of virtual markers with at least one matching ID was above 99% in all cases, and Tgt match did not exceed this value in all scenes (S3H Fig).

Thus, under this condition, vmT-DLC outperformed maDLC in terms of Tgt match. Furthermore, vmT-DLC demonstrated the ability to achieve tracking accuracy beyond the precision of the virtual markers. However, both these observations were limited to the OC scenes because they were not observed in the nOC scenes. Notably, compared with that of the five-mice tracking experiment, the accuracy of SLEAP was the lowest. Consequently, while vmT-LEAP also showed higher accuracy than SLEAP, it did not perform better than maDLC or vmT-DLC.

### Relationship between annotation frame count and tracking accuracy

vmT-DLC showed a high Tgt match regardless of the contrast with the background color or the presence of occlusion and crowding. Another notable finding was that vmT-DLC produced no FNs. By narrowing the match criteria in maDLC to only predict keypoints, the difference in accuracy between maDLC and vmT-DLC was reduced. Therefore, we examined whether repeated training with maDLC could reduce FNs and potentially achieve a Tgt match comparable to that of vmT-DLC by analyzing the relationship between annotation frame count and accuracy. The points plotted for each annotation frame count represent the number of iterations (retraining after refinement) in maDLC, with the leftmost point corresponding to iteration-0.

Focusing on the OC scenes in the three-mice tracking experiment, we first compared the changes in Tgt match, Pred match, and FNs across iterations (Friedman test with Bonferroni correction, n = 12). For maDLC, although significant differences were observed at some points between iteration-0 (400 frames) or iteration-1 (650 frames) and other time points, there were no significant differences between iteration-1 and iteration-11 (3886 frames) across all three evaluated metrics (Fig 3G). Similarly, for vmT-DLC, while significant differences were observed between iteration-0 (200 frames) and iteration-1 (400 frames) and subsequent iterations, there were no significant differences after iteration-2 (598 frames) (Fig 3G).

Next, we compared Tgt match and Pred match at each iteration with the median Tgt match results of vmT-DLC (equivalent to Pred match) (pairwise comparisons with Bonferroni correction). Tgt match in maDLC remained consistently and significantly lower at all iterations (Fig 3G). However, Pred match in maDLC showed no significant differences across iterations (Fig 3G).

In summary, under these conditions, increasing the annotation frame count in maDLC had minimal impact on the number of FNs, and further refinement with additional frames offered limited improvement in Tgt match or Pred match.

### Application of tracking data to Keypoint-MoSeq

The posture analysis tool Keypoint-MoSeq [18] was applied to data obtained from maDLC and vmT-DLC in a three-mice tracking experiment with six keypoints. In this study, we aimed to evaluate whether differences in tracking accuracy affected the results of posture analysis. To this end, we first investigated the similarity between data obtained from different tracking methods.

Additionally, in scenes where discrepancies occurred, we examined whether the results derived from the more accurate vmTracking provided a more precise model—specifically, one that extracted features more consistent with the actual behavior.

After model creation, we analyzed the tracking data from maDLC and vmT-DLC. The results shown are based on a single video analyzed with both maDLC and vmT-DLC. The similarity dendrogram indicates that nearly identical syllable patterns were obtained using both methods (S4A Fig). Cross-referencing the similarity dendrogram results with the syllable plots, syllables 6, 8, and 11 were identified as low-movement or near-stationary states, and they were consolidated for further evaluation (S4B Fig). To assess the similarity of these syllables identified by maDLC and vmT-DLC, we examined which syllable each posture (behavior) was assigned to within the same frames and summarized the results in a confusion matrix (S4C Fig). Frames where the same syllables were assigned in both maDLC and vmT-DLC had a higher proportion compared to those with different assignments. Cohen’s kappa coefficient, an index of agreement, was 0.688, which is generally considered a substantial level of agreement (S4C Fig). From the confusion matrix, the proportion of syllables identified differently by maDLC and vmT-DLC was generally below 1%, except for three cases where the proportion exceeded 1%: syllables identified as IV in maDLC but VI in vmT-DLC, V in maDLC but VI in vmT-DLC, and VI in maDLC but IV in vmT-DLC (cells shaded in purple) (S4C Fig). As “Others” included minor syllables not represented in the similarity dendrogram or syllable plots, we excluded patterns involving “Others” from this evaluation.

In S4D Fig, we verified the validity of syllable assignments by examining tracking images in scenes where these relatively frequent mismatched patterns continued for 2 s (60 frames) or more. The upper row shows an example of the pattern where maDLC identified syllable IV and vmT-DLC identified VI (here, syllable 6), with magenta-outlined frames highlighting instances of these assignments. The rightward turn observed from 0 to 1 s matched the pattern of maDLC’s syllable IV.

However, syllable IV displayed a leftward body orientation at the start of the movement, which did not align in this aspect. In contrast, vmT-DLC’s syllable VI did not capture the rightward turning motion but did align in terms of a rightward body orientation and matched the mostly stationary pattern seen after 2 s (S4D Fig, upper row). The lower row illustrates an example in which maDLC identified syllable VI (here, syllable 6) and vmT-DLC identified syllable IV. From −1 to 0 s, a rightward turn was observed, which appeared undetected in maDLC’s syllable 6; however, the immobility following 1 s seemed consistent. vmT-DLC’s syllable IV appeared to capture the movement from −1 to 0 s. Although this syllable represents a rightward turn pattern, the movement range is minimal, showing similarity with the stationary pattern after 1 s (S4D Fig, lower row).

In summary, substantial agreement was observed between the syllables identified by maDLC and vmT-DLC. For scenes where the identified syllables were inconsistent, cross-referencing them with the actual observed behavior revealed that both maDLC and vmT-DLC sometimes identified a scene with movement as a non-movement syllable. On the contrary, the posture (e.g., left-curved, straight, or right-curved) extracted by vmT-DLC matched the actual posture, whereas there were cases where the posture extracted by maDLC slightly differed from the actual behavior.

### Tracking an object entering and exiting from a video frame

A validation study was conducted to simulate experiments in which individuals enter and exit the video frame. Typically, maDLC does not detect markers of individuals not present in the frame, whereas saDLC outputs all markers regardless of individual presence. This setup explored the impact on tracking when individuals entered and exited the recording frame in vmT-DLC (S5 Fig). A simplified flow of the experiment is shown in S5A Fig. We manually removed and introduced one mouse at a time in conditions featuring three mice against a white backgroun. During annotation, stray keypoints of mice removed from the frame were labeled on a remaining mouse (S5B Fig).

Accurate tracking was conducted with all three mice present, according to the virtual markers (S5C Fig, lower panel for the three-mice condition). However, when removing one mouse from the arena, the stray keypoints that lost their tracking target generally detected another remaining mouse, according to the annotations (S5C Fig, lower panel for the two-mice condition). Similarly, removing another mouse generally caused stray keypoints to detect the single remaining mouse (S5C Fig, lower panel of single-mouse conditions). When reintroducing the mice into the arena, tracking resumed with the correct IDs according to the virtual markers (S5C Fig, lower panel for the transitions from one mouse to two- and three-mice conditions).

For quantitative evaluation, we calculated the distances between identical keypoints of the three mice in each frame and evaluated their distribution (S5D Fig). We defined overlap as a distance of less than 10 pixels between identical keypoints from two different IDs and calculated the frequency of frames in which such overlap occurred for each keypoint. The average percentage of frames in which stray keypoints overlapped with identical keypoints in another mouse was greater than 90% for the six keypoints (S5D Fig and S1 and S2 Table). The high frequency of overlap for ID2 and ID3 in the two-mice condition and for the three IDs in the one-mouse condition indicate that in vmT-DLC, stray keypoints that lost their original tracking targets frequently aligned with the identical keypoints of remaining mice.

These overlapping keypoints are unnecessary in the finalized tracking results. To address this aspect, we developed a Python code (https://doi.org/10.5281/zenodo.14249295) to remove specified time ranges and keypoints (S10C Fig) and applied it to these data (S5Ea–c Fig and S1 Video). This code also proved effective in maDLC, removing unnecessary ID data in cases of overlapping IDs, thereby enabling the creation of accurate virtual marker videos with colors corresponding to the correct ID (S5Ed Fig).

### Physical marker tracking

In vmTracking, virtual markers were used as substitutes for physical markers, and they showed particularly high accuracy in vmT-DLC. However, whether virtual markers could truly replace physical markers remained unclear because unforeseen factors may have facilitated vmTracking. We performed a comparison by tracking physical marker videos using maDLC and saDLC (Fig 4 and S5 Video). To this end, we tracked more lateralized body parts than the previous six points. In doing so, we used the same annotated data across all conditions to ensure consistency.

**Fig 4.**
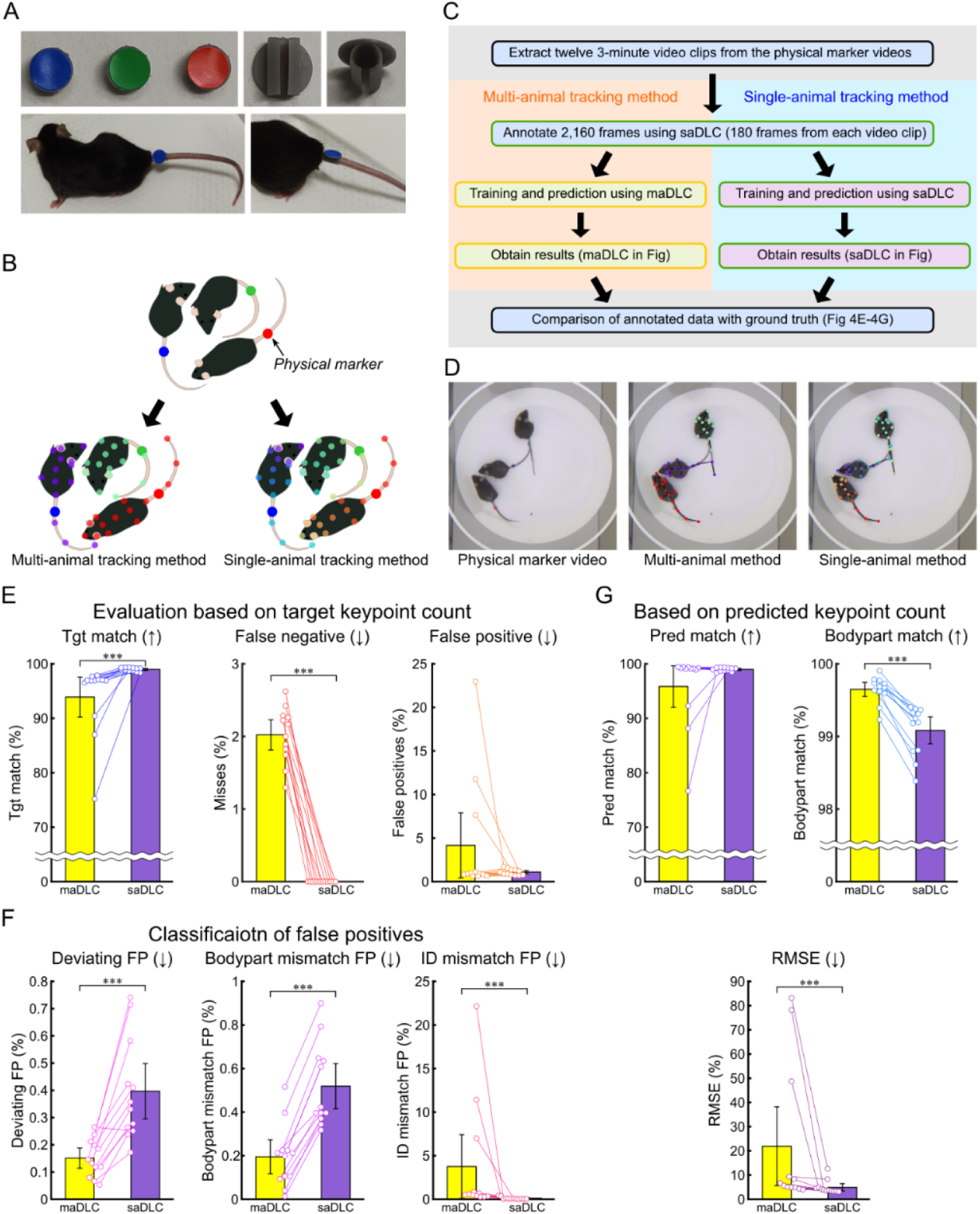
Multi-animal tracking with physical markers. (A) Physical markers. Physical marker attachments were created using a 3D printer, with colored tack stickers added as visual markers, and attached to the mouse’s tail. (B) Workflow of this experiment. The same annotation data were used to track a multi-animal video with physical markers, using both multi-animal DeepLabCut (maDLC) and single-animal DeepLabCut (saDLC). The annotation data served as the ground truth for accuracy evaluation. (C) Schematic of annotations in maDLC and saDLC. The yellow box in the schematic represents the processes using maDLC, while the green box represents the processes using saDLC. (D-F) Various percentage-based metrics were compared using the annotated data as ground truth (GT). The comparisons included maDLC and saDLC. (D) Snapshots of the physical marker video and the predictions from maDLC and saDLC. (E) The evaluation was conducted as a percentage of the target keypoints, with the total number of GT keypoints as the denominator. The percentages of matches (Tgt match), false negatives, and false positives were calculated based on the number of GT keypoints. (F) False positives from (E) were classified into deviations exceeding the threshold from all GT (Deviating FP), mismatches in predicted body parts (Bodypart mismatch FP), and cases where the predicted body part was correct but the ID was incorrect (ID mismatch FP). (G) Evaluation was conducted as a percentage of the predicted keypoints, excluding FNs, using the total predicted keypoints as the denominator, and the metrics included matches (Pred match), body part matches (Bodypart match), irrespective of ID, and root mean square error (RMSE). Arrows indicate whether higher or lower values are better. Data are shown as mean ± 95% confidence interval (n = 12 scenes). Statistical analysis was performed using the Wilcoxon sign-ranked test (n = 12), ***: p < 0.001.

The physical markers were designed to avoid creating noticeable visual contrasts, such as differences in body color, aiming to offer similar features to virtual markers. The physical markers were created using a 3D printer, designed to be attached by threading onto the tail. Colored tack stickers were applied to the markers, allowing individual mice to be identified by color (Fig 4A). This setup enabled consistent annotations across frames by using marker color as a cue, similar to when using virtual markers (Fig 4B). A brief outline of the experiment is provided in Fig 4C, and examples of the physical marker images along with tracking results from maDLC and saDLC are shown in Fig 4D.

A comparison based on the number of target points (Friedman test with Bonferroni correction, n = 12) revealed significantly better performance for saDLC than maDLC in Tgt match (p < 0.001) and FNs (p < 0.001) (Fig 4E). While there was no significant difference in FPs, within the FP classifications, saDLC performed significantly better only in ID mismatch FPs (p < 0.001) (Fig 4E and 4F). Conversely, maDLC was significantly better than saDLC in Deviating FPs (p < 0.001) and Bodypart mismatch FPs (p < 0.001) (Fig 4F). An evaluation based on predicted keypoints revealed significantly better performance for saDLC in terms of RMSE (p < 0.001), compared with maDLC—which performed significantly better in Bodypart match (p < 0.001) (Fig 4G). No significant difference was observed in Pred match (Fig 4G).

Thus, both maDLC and saDLC were used to track videos with physical markers, resulting in similar outcomes as vmTracking. Specifically, saDLC showed better performance than maDLC in Tgt match, FNs, and ID mismatch FPs but worse performance in Deviating FPs and Bodypart mismatch FPs.

### Tracking of black mice against white background using 14 lateralized keypoints

In this validation, in addition to tracking previously used markerless videos and virtual marker videos, we also introduced tracking of another virtual marker videos created based on results from idtracker.ai [19]. Additionally, for markerless video tracking with SLEAP, we examined both the top-down and bottom-up approaches as analysis pipelines. Fig 5A and 5B shows a simplified workflow of the experiment, and Fig 5C and S6 Video presents an example of the tracking results.

**Fig 5.**
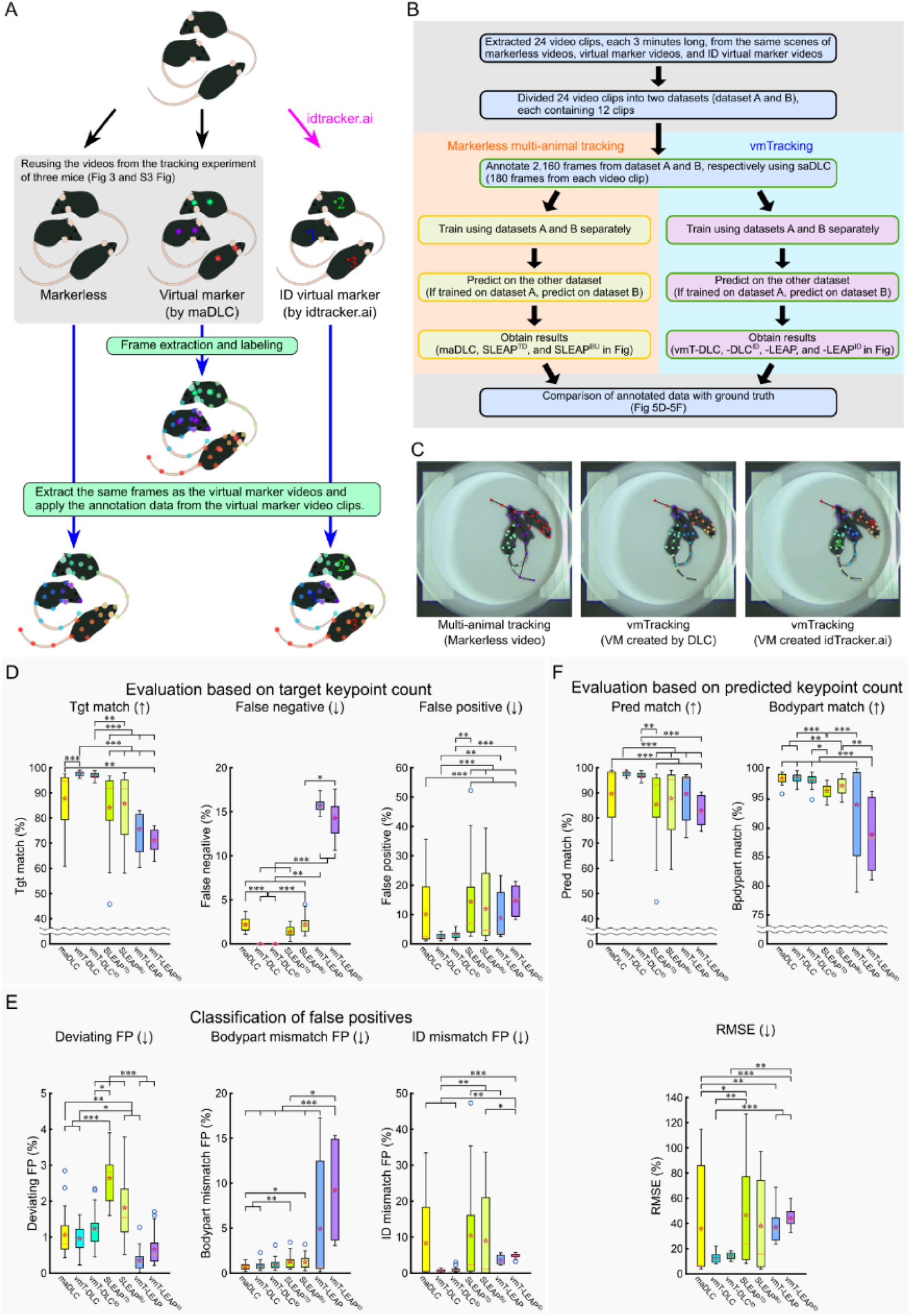
Validation of vmTracking under various conditions with a common annotation dataset. (A) Process of creating a common annotation dataset. In this verification, to create a common annotation dataset, frames were extracted from the virtual marker video, and 14-point lateralized labeling was applied. This annotated data were then applied to the same frames extracted from both the markerless video and the ID virtual marker video created by idtracker.ai. (B) Flow of this verification. The yellow box in the schematic represents the processes using multi-animal tracking tools, while the green box represents the processes using single-animal tracking tools. (C) Example snapshot predicted by DeepLabCut (DLC) from the markerless video, virtual marker video created by DLC, and ID virtual marker video. (D–F) Various percentage-based metrics were compared using the annotated data as ground truth (GT). The methods compared included multi-animal DLC (maDLC), virtual marker tracking with single-animal DLC (vmT-DLC), vmT-DLC with ID virtual marker videos (vmT-DLC^ID^), SLEAP’s multi-animal top-down method (SLEAP^TD^), multi-animal bottom-up method (SLEAP^BU^), virtual marker tracking with SLEAP (vmT-LEAP), and vmT-LEAP with ID virtual marker videos (vmT-LEAP^ID^). (D) Evaluation was conducted as a percentage of the target keypoints, with the total number of GT keypoints as the denominator. The percentages of matches (Tgt match), false negatives, and false positives were calculated based on the number of GT keypoints. (E) False positives from (D) were classified into deviations exceeding the threshold from all GT (Deviating FP), mismatches in predicted body parts (Bodypart mismatch FP), and cases where the predicted body part was correct but the ID was incorrect (ID mismatch FP). (F) Evaluation was conducted as a percentage of the predicted keypoints, excluding FNs, using the total predicted keypoints as the denominator, and the metrics included matches (Pred match), body part matches (Bodypart match), irrespective of ID, and root mean square error (RMSE). Arrows indicate whether higher or lower values are better. The results are presented as box plots, where the bottom of the box represents the first quartile (Q1), the top represents the third quartile (Q3), the orange line inside the box indicates the median, the whiskers extend to 1.5 times the interquartile range (Q3-Q1), blue circular markers denote outliers, and hollow red star markers represent the mean. Statistical analysis was performed using the Friedman test with Bonferroni correction (n = 24). *: p < 0.05, **: p < 0.01, ***: p < 0.001.

Here, we retained the designations “maDLC,” “vmT-DLC,” and “vmT-LEAP” as previously used. For markerless multi-animal tracking with SLEAP, we denote the top-down approach as “SLEAP^TD^” and the bottom-up approach as “SLEAP^BU^.” Additionally, for tracking virtual marker videos created with idtracker.ai, we labeled them as “vmT-DLC^ID^” and “vmT-LEAP^ID^,” depending on the tool used.

A comparison of metrics based on the number of target keypoints (Friedman test with Bonferroni correction, n = 24) revealed that vmT-DLC outperformed maDLC in Tgt match (p < 0.001) and FNs (p < 0.001). However, no significant difference was observed for FPs or the classified FPs (Fig 5D and 5E). For metrics based on predicted keypoints, no significant differences were observed between vmT-DLC and maDLC across all metrics, mirroring the trend observed in the six-keypoint tracking (Fig 5F). A comparison of vmT-DLC^ID^ and maDLC, vmT-DLC^ID^ showed better results only in FNs (p < 0.001), with no significant differences in other metrics (Fig 5D–5F).

Notably, no significant differences were found between vmT-DLC and vmT-DLC^ID^ across all metrics (Fig 5D–5F), indicating that vmT-DLC and vmT-DLC^ID^ performed statistically equivalently.

In contrast, for vmT-LEAP and vmT-LEAP^ID^, the only metric with significantly better results compared to maDLC, vmT-DLC, vmT-DLC^ID^, SLEAP^TD^, and/or SLEAP^BU^ was Deviating FPs (Fig 5D–5F). vmT-LEAP performed significantly better than maDLC (p < 0.01), vmT-DLC^ID^ (p < 0.001), SLEAP^TD^ (p < 0.001), and SLEAP^BU^ (p < 0.001) (Fig 5E). vmT-LEAP^ID^ performed significantly better than vmT-DLC^ID^ (p < 0.001), SLEAP^TD^ (p < 0.001), and SLEAP^BU^ (p < 0.001) (Fig 5E). Similar to DLC, there were no significant differences between vmT-LEAP and vmT-LEAP^ID^ across all metrics (Fig 5D–5F), showing statistically equivalent. Additionally, according to the video, vmT-LEAP had difficulty tracking the tail in most cases (S6 Video).

These findings suggest that vmT-DLC enables consistently high match rates across different keypoint tracking conditions, and that virtual marker videos created with idtracker.ai can achieve accuracy comparable to those created with DLC. In contrast, for this condition, vmT-LEAP resulted in lower performance, compared with markerless video tracking.

### Evaluation of centroid tracking

To date, evaluations have focused on individual keypoints. However, by using the centroid derived from tracked keypoints as a representative value, we aimed to assess the overall tracking accuracy. This approach evaluates how accurately the entire mouse body was tracked, mitigating the impact of outlier keypoints. Specifically, in addition to the centroid based on 11 keypoints representing the torso region from the 14-point lateralized tracking (obtained with maDLC, vmT-DLC, SLEAP^BU^, and vmT-LEAP), we evaluated the centroid obtained from idtracker.ai (ID). SLEAP^BU^ was used instead of SLEAP^TD^ because no outliers were observed in the match evaluation in Fig 5. Given that centroids calculated from posture-tracking keypoints may differ in characteristics from those obtained from idtracker.ai, we defined a match as when a centroid was within 20 pixels of any of the 3 body-axis keypoints in the GT data (Fig 6A). Using this criterion, we evaluated whether tracking was centered around the area of the correct mouse’s torso.

**Fig 6.**
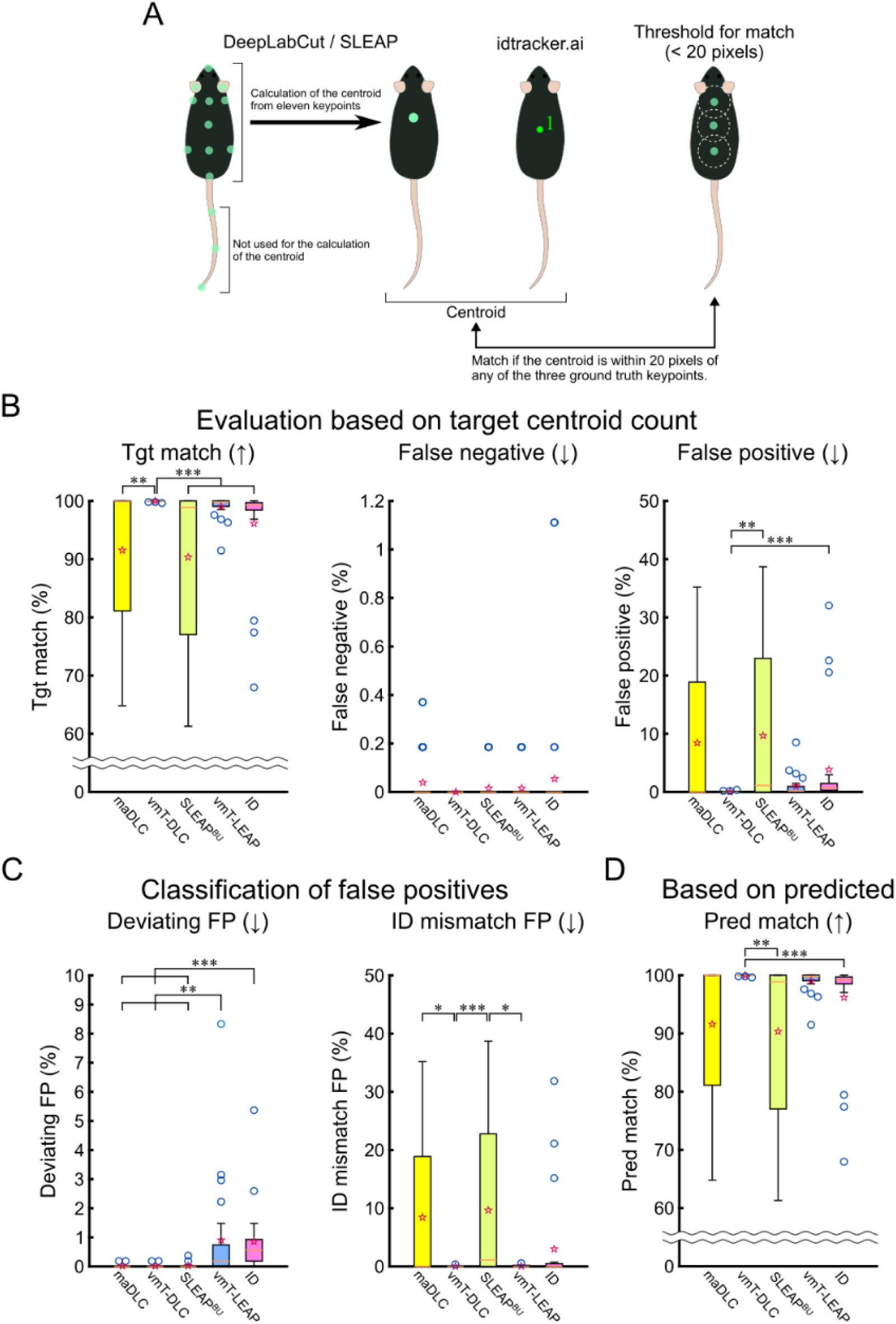
Comparison of tracking accuracy based on centroids. (A) Method for calculating and evaluating centroids. The centroids obtained from DeepLabCut and SLEAP were calculated based on 11 predicted keypoints of 14, excluding the tail. The centroid from idtracker.ai was used as is. These centroids were evaluated based on whether they were within 20 pixels of any of the 3 ground truth points along the body axis, to assess whether the centroids were positioned on the mouse’s body. This evaluation was performed for the centroid using multi-animal DeepLabCut (maDLC) and virtual marker tracking with single-animal DeepLabCut (vmT-DLC), SLEAP’s multi-animal bottom-up method (SLEAP^BU^) and virtual marker tracking with SLEAP (vmT-LEAP), and idtracker.ai (ID). (B) The evaluation was conducted as a percentage of the target keypoints, with the total number of GT keypoints as the denominator. The percentages of matches (Tgt match), false negatives, and false positives were calculated based on centroids. (C) The false positives from (B) were classified into deviations exceeding the threshold from all GT (Deviating FP) and cases where the predicted ID was incorrect (ID mismatch FP). (D) Matches (Pred match) were evaluated as a percentage of the total number of predicted centroids, excluding false negatives. Arrows indicate whether higher or lower values are better. The results are presented as box plots, where the bottom of the box represents the first quartile (Q1), the top represents the third quartile (Q3), the orange line inside the box indicates the median, the whiskers extend to 1.5 times the interquartile range (Q3-Q1), blue circular markers denote outliers, and hollow red star markers represent the mean. Statistical analysis was performed using the Friedman test with Bonferroni correction (n = 24). *: p < 0.05, **: p < 0.01, ***: p < 0.001.

A comparison of metrics based on the number of target keypoints (Friedman test with Bonferroni correction, n = 24) revealed that vmT-DLC achieved a significantly higher Tgt match than did maDLC (p < 0.01), SLEAP^BU^ (p < 0.001), and ID (p < 0.001), while no significant difference was observed compared with vmT-LEAP (Fig 6B). No significant differences were found in FNs across methods (Fig 6B). For FPs, vmT-DLC showed significantly better results than SLEAPBU (p < 0.01) and ID (p < 0.001), but no significant differences were found compared with maDLC and vmT-LEAP (Fig 6B). In the FPs classification, maDLC (p < 0.01), vmT-DLC (p < 0.01), and SLEAP^BU^ (p < 0.01) performed significantly better than vmT-LEAP, and maDLC (p < 0.001), vmT-DLC (p < 0.001), and SLEAP^BU^ (p < 0.001) showed better performance than ID (Fig 6C). For ID match FPs, vmT-DLC showed significantly better performance than did maDLC (p < 0.05) and SLEAP^BU^ (p < 0.001), while vmT-LEAP performed significantly better than SLEAP^BU^ (p < 0.05). No significant differences were observed with ID (Fig 6C). Based on metrics from predicted keypoints, vmT-DLC showed significantly better results than SLEAP^BU^ (p < 0.01) and ID (p < 0.001), but no significant difference was observed compared with maDLC (Fig 6D).

Thus, vmT-DLC demonstrated a high Tgt match and performed well in terms of FPs. Notably, while vmT-LEAP showed lower performance than vmT-DLC in keypoint-based evaluation, the two methods showed comparable performance in centroid evaluation.

### Verification of applicability to fish

The applicability of vmTracking to other animal species was validated by attempting to track a school of 10 fish. Fig 7A shows an overview of the experimental setup. In this study, the validation was conducted using a single video consisting of 2,259 frames. Fig 7B is a snapshot of the tracking video, where 10 fish swim within a rectangular tank. For each fish, we tracked five equidistant points along the body axis, using the second and fourth keypoints as virtual markers (Fig 7C).

**Fig 7.**
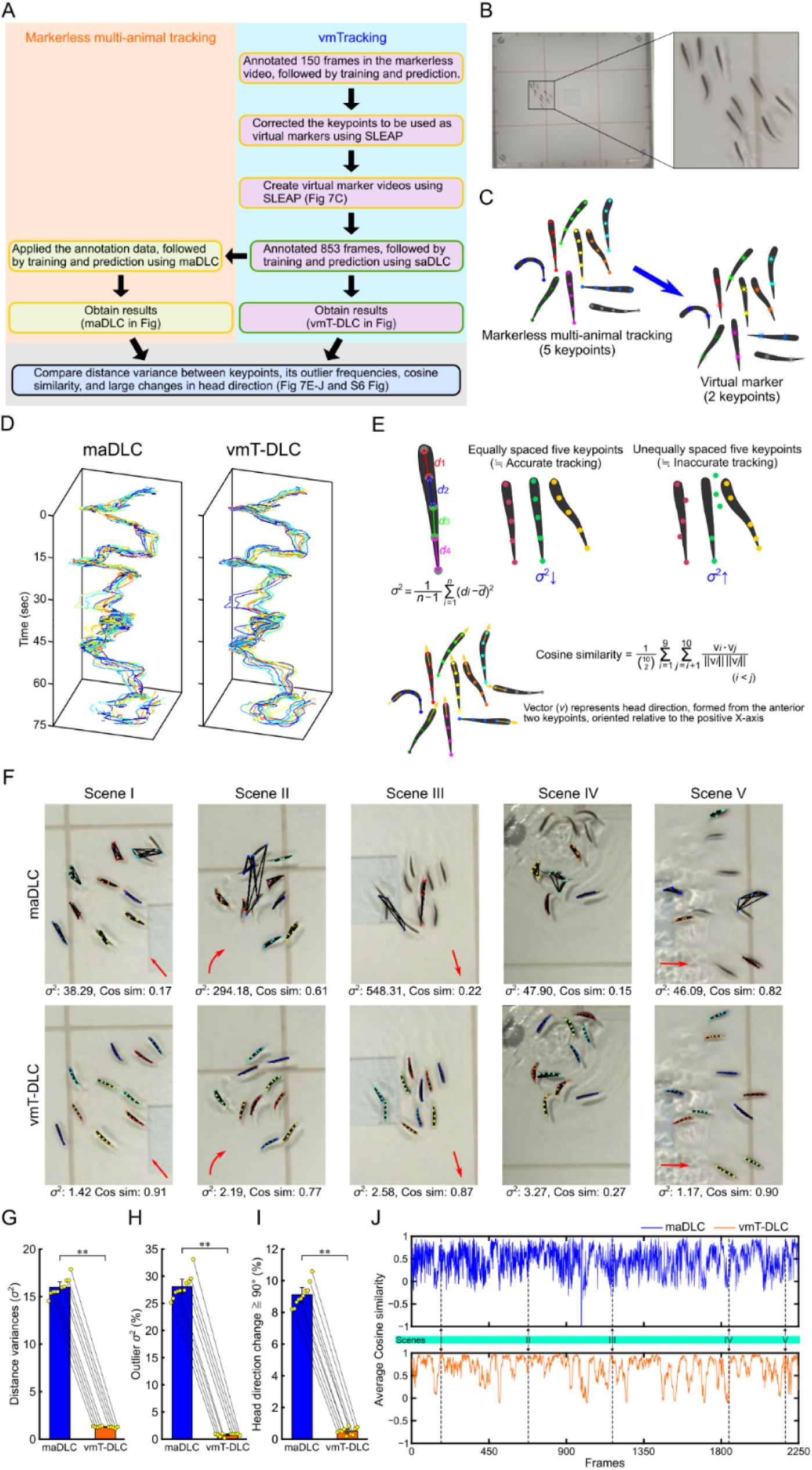
Comparison of markerless multi-animal pose tracking and vmTracking in multi-fish tracking. (A) Overview of the verification process. The yellow box in the schematic represents the processes using multi-animal tracking tools, while the green box represents the processes using single-animal tracking tools. In this study, the evaluation was conducted on a single video consisting of 2,259 frames. (B) Image of the tracked school of fish. Right panel: enlarged view of the rectangle in the left panel. (C) Schematic of virtual marker creation. Five keypoints per fish were tracked, with two designated as virtual markers. (D) Swimming trajectories of 10 fish based on the foremost (head) keypoint for multi-animal DeepLabCut (maDLC) and virtual marker tracking with single-animal DeepLabCut (vmT-DLC). (E) Methods for evaluating tracking. Based on tracking with equidistant keypoints, variance in distance between adjacent keypoints was used as a basic indicator of accurate tracking (larger variance indicates lower accuracy) (Upper row). Cosine similarity (Cos sim) of head direction for 10 fish was calculated using angles formed by the 2 foremost keypoints (Bottom row). (F) Example photos from five scenes and corresponding values for variance ( ^2^) and Cos sim. Red arrows in the photos indicate general movement direction of the school of fish. Scene IV did not show a clear direction of movement as a group (not indicated). (G–I) Median variance for each fish (plot). (G), the number of frames showing outliers in variance for each fish based on the interquartile range method (plot); (H), the number of frames for each fish where the head direction changed by more than 90° between frames (approximately 0.03 s) (plot) (I), with bars and error bars representing mean ± 95% confidence interval calculated from the medians of all 10s fish (n= 10). Statistical analysis was performed using the Wilcoxon signed-rank test (n = 10), **: p < 0.01. (J) Time series changes in Cos sim. Scenes I-V correspond to Scenes I-V in (F).

Virtual marker videos were created using SLEAP, and the tracking comparison between maDLC and vmT-DLC was conducted (S1 and S7 Video). The trajectories for keypoint 1 (head) in both maDLC and vmT-DLC were demonstrated (Fig 7D). While maDLC showed intermittent trajectories, vmT-DLC consistently produced smooth trajectories.

Accuracy was verified based on equidistant tracking along the body axis, using the variance of the distances between adjacent keypoints, where higher values reflect lower accuracy (Fig 7E upper row). Additionally, to detect schooling—a behavior characteristic of fish schools involving swimming in the same direction—the head direction of each fish was calculated, and their cosine similarity value (Cos sim) was measured based on the orientation of the two anterior keypoints (Fig 7E lower row). Fig 7F displays snapshots of five exemplary scenes, along with their respective variance ( ^2^) and Cos sim values, demonstrating that vmT-DLC more accurately tracked each fish. The comparison of median variance for each fish revealed significantly lower variance in vmT-DLC (p < 0.01, Wilcoxon signed-rank test, n = 10, Fig 7G). Analysis also revealed significantly fewer frames with outlier variances in vmT-DLC when counting frames where each fish’s variance values were outliers (p < 0.01, Fig 7H). These results indicate that vmT-DLC more effectively detected keypoints at consistent intervals, essential for accurate tracking. In comparisons based on head direction, the frequency of head direction changes of more than 90° between frames (approximately 0.03 s) was significantly lower in vmT-DLC (p < 0.01, Fig 7I). The time series variation of Cos sim for both maDLC and vmT-DLC appeared to follow similar patterns; however, maDLC exhibited greater fine-scale variability, resembling noise (Fig 7J), indicating more frequent changes in head direction similarity between frames. Furthermore, in an analysis using the GT consisting of 410 frames, a correlation with GT Cos sim revealed a stronger correlation between GT and vmT-DLC (*ρ* = 0.967, n = 407, S6A Fig right) compared to GT and maDLC (*ρ* = 0.485, n = 407, S6A Fig left). This indicates that the results from vmT-DLC aligned closely with those of GT. We also calculated the same GT-based metrics as those evaluated in mouse tracking and examined their respective proportions (S6B and S6C Fig). In vmT-DLC, the Tgt match rate and Pred match rate exceeded 99%, indicating highly accurate tracking. These results demonstrated that vmT-DLC also achieved greater accuracy than maDLC in tracking fish schools.

### Tracking of coordinated poses and movements among dancers

vmTracking was further applied to humans. Fig 8A shows an overview of the experimental setup. 7 dancers were tracked using 19 keypoints, from which, videos were created with 2 virtual marker configurations: one with virtual markers on the head and trunk, and another with six virtual markers, including the head, trunk, both elbows, and both knees (Fig 8B). The snapshots on the right display examples of the markerless video, the two-point virtual marker video, and the six-point virtual marker video. Tracking was conducted using markerless or virtual marker tracking in these videos. Reviewing these tracking videos revealed inaccuracies in keypoint predictions in both maDLC and vmT-DLC. While maDLC occasionally switched the entire set of keypoints to a different dancer, no such phenomenon was observed with vmT-DLC (S8 Video). By focusing on scenes with similar movements, we evaluated the accuracy of detecting the coordinated movements of the seven dancers over a 2-s period.

**Fig 8.**
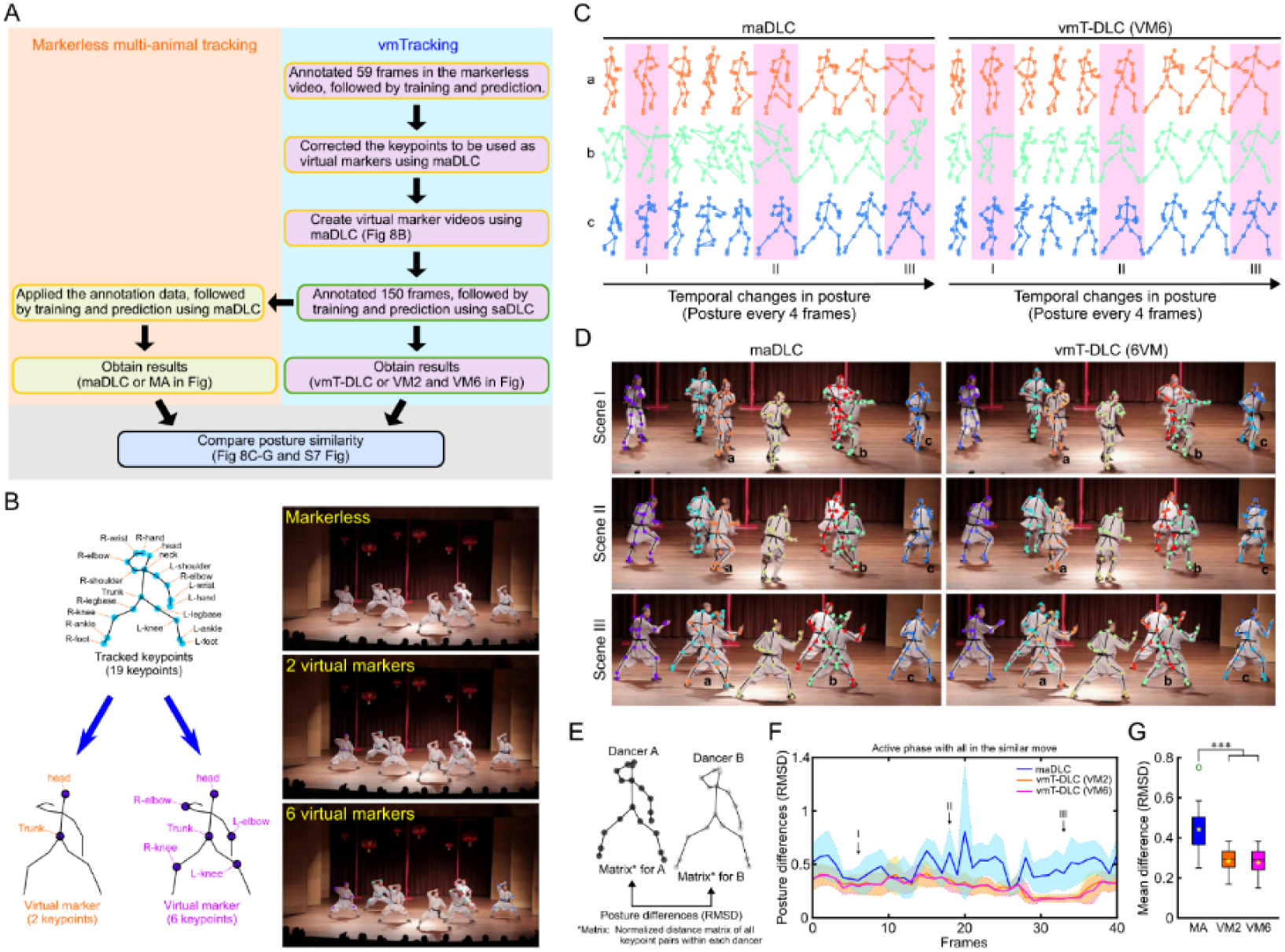
Comparison of markerless multi-animal pose tracking and vmTracking in multiple dancer tracking. (A) Overview of the verification process. The yellow box in the schematic represents the processes using multi-animal tracking tools, while the green box represents the processes using single-animal tracking tools. (B) Nineteen keypoints were tracked, and two or six keypoints were designated as virtual markers. The snapshots on the right are examples of markerless and virtual marker videos. (C) Extraction of tracking data for three dancers from a scene where seven dancers perform similar movements over 2 s, displayed at nine points every four frames. The results show multi-animal DeepLabCut (maDLC) and virtual marker tracking with single-animal DeepLabCut (vmT-DLC) with six-point virtual marker video (VM6). The dancers labeled a–c correspond to dancers a to c in (D), and I to III correspond to I to III in (D) and (F). (D) Comparison of actual tracking photos for scenes I to III shown in (C). (E) Method for assessing the similarity of poses among the seven dancers. A standardized distance matrix was created for each dancer, comparing all keypoint pairs, and the root mean square deviation (RMSD) of these matrices was calculated between dancers. (F) Time series variation in the average RMSD values for maDLC, vmT-DLC with two-point virtual marker video (VM2), and VM6 of all dancer pairs for each frame over 2 s, with the shaded area indicating standard deviation. (G) Box plots of RMSD values for maDLC (MA), VM2, and VM6, calculated for all frames over a 2-s period, are presented. The bottom of the box represents the first quartile (Q1), the top represents the third quartile (Q3), the line inside the box indicates the median, the whiskers extend to 1.5 times the interquartile range (Q3-Q1), green circular markers denote outliers, and yellow star markers represent the mean. Statistical analysis was performed using the Friedman test with Bonferroni correction (n = 41). *: p < 0.05, **: p < 0.01, ***: p < 0.001.

Fig 8C shows representative examples of the temporal changes in the poses of three dancers tracked by maDLC and vmT-DLC. In maDLC, keypoints occasionally appeared in positions distant from their correct locations, considerably more than that in vmT-DLC. Actual images from these scenes were displayed to illustrate the real tracking images (Fig 8D). To quantitatively assess the similarity of the poses of the seven dancers, a normalized distance matrix for all keypoint pairs of each dancer was created, and the root mean square deviation (RMSD) between the dancers was compared (lower RMSD indicates higher similarity) (Fig 8E). vmT-DLC consistently showed RMSDs of 0.5 or lower, whereas maDLC sporadically displayed spikes in higher values, indicating lower pose similarity in some frames (Fig 8F). Overall, vmT-DLC demonstrated significantly lower RMSD values (p < 0.001, Bonferroni correction with Friedman test, n = 41, Fig 8G). Similarly, the analysis evaluated scenes where the seven dancers maintained similar poses while stationary (S7 Fig). Even in these stationary scenes, maDLC showed keypoint switches, especially among overlapping dancers, whereas vmT-DLC achieved stable tracking (S7A and S7B Fig), and showed significantly lower average RMSD values (p < 0.001, Bonferroni correction with Friedman test, n = 41, S7C Fig). We also calculated the same GT-based metrics as those evaluated in mouse tracking and examined their respective proportions (S7D and S7E Fig). The GT consisted of 150 frames (out of a total video length of 703 frames). In vmT-DLC, the Tgt match rate and Pred match rate exceeded 99%, indicating highly accurate tracking.

## Discussion

Despite advancements in markerless multi-animal pose tracking tools, overcoming occlusion and crowding remains a challenge. Studies using saDLC to simultaneously track one black C57BL/6N mouse and one white CD1 mouse have been reported [17], suggesting that tracking multiple animals with saDLC is feasible when there is a visual difference. Although attaching visually identifiable physical markers to animals could be a potential solution, physical markers may unpredictably affect behavior. In this study, we adopted a straightforward strategy of applying virtual markers to videos of markerless animals, allowing us to distinguish between animals without the need for physical markers. By tracking multiple animals with virtual markers using a single-animal pose tracking tool, we improved the accuracy of traditional multi-animal pose tracking in scenarios involving occlusion and crowding. The approach of locating individuals first and then tracking them was also adopted in SLEAP’s top-down method [8]. However, vmTracking achieved very high accuracy by incorporating human intervention to reliably assign individual identification information and then using that information for tracking. In this section, we discuss the advantages and limitations of this method, namely vmTracking, as well as its implications for future research directions and its impact on this field.

### vmTracking methodological enhancements and efficiency

vmTracking showed superior performance compared to traditional markerless multi-animal pose tracking methods, consistently improving accuracy across various conditions, including tracking three or five animals and managing occlusion and crowding. Notably, vmT-DLC improved keypoint matching rates in scenes where traditional markerless multi-animal tracking methods such as maDLC and SLEAP previously showed less than 50% matching accuracy, increasing it to approximately 90%. Conversely, virtual marker tracking using SLEAP’s single-animal tracking approach (vmT-LEAP) displayed limited effectiveness, with performance depending on the specific tracking target and video conditions. In particular, the performance for tracking the 14-point lateralized keypoints was poor, which is likely due to difficulties in tracking the tail. As such, using saDLC in the virtual marker tracking step is considered optimal.

The high accuracy achieved by vmT-DLC appears to be linked to its ability to ensure that all keypoints are consistently predicted, thereby preventing FNs in the tracking process. While this may cause a slight increase in FPs, the overall high accuracy suggests that most predictions are valid. Consequently, vmT-DLC significantly mitigated issues such as missed detections, which are common in conventional markerless multi-animal tracking. Another factor contributing to the high accuracy is the drastic reduction in ID switches and ID mismatches that commonly occur in scenes with occlusion and crowding. These results confirm that vmT-DLC represents a robust solution for addressing occlusion and crowding.

vmTracking requires a two-step process. First, create virtual markers. Second, track with saDLC. Therefore, the high accuracy of vmTracking could also be interpreted as the result of inheriting the corrected maDLC results during virtual marker creation and further training with saDLC. However, increasing the number of training frames with maDLC or SLEAP does not necessarily achieve the accuracy levels reached by vmT-DLC. With maDLC, even when increasing the number of annotation frames, FNs (false negatives) persist at a certain rate without much reduction, posing an obstacle to improving accuracy. This may partly be due to the limitation in DLC version 2.2.3 used in this study, which cannot supplement FN keypoints during refinement annotation. Even if FNs could be corrected in annotated frames, this would be possible in only a subset of frames, suggesting a limited impact on reducing FNs overall. Ultimately, unless FNs can be reduced, it would be impossible to reach the accuracy levels achieved by vmT-DLC in scenes with occlusion or crowding.

Another argument against vmTracking might be that the effort spent on correcting data to create virtual markers could simply be used for correcting the results of markerless tracking directly to obtain accurate tracking outcomes. However, notably, the presence of virtual markers with incorrect IDs or missing markers seemed to have little to no impact on reducing tracking accuracy. It can be assumed that the cost of modification work to create virtual markers is overwhelmingly smaller than that of modification work required to obtain correct tracking results. While there may be a threshold for the acceptable frequency of these less-than-ideal virtual markers, this study suggests that even in scenes with occlusion or crowding, vmT-DLC could likely achieve 90% or more accuracy if 70–80% of the virtual markers were assigned correct IDs. This 80% accuracy suggests that even if 10% FNs occurred in markerless tracking, achieving 90% accuracy with the remaining data would suffice, aligning with the accuracy of unmodified maDLC and SLEAP. Although some researchers may consider this level of accuracy sufficient for practical tracking while others may not, it is adequate for creating virtual markers.

Thus, maDLC can achieve high accuracy with minimal annotation frames, even in scenes with occlusion or crowding, and this likely applies to SLEAP as well. This high capability will greatly assist in virtual marker creation by minimizing the training and manual corrections required for creating virtual markers. The requirement for further training and thorough correction to create more accurate virtual markers is considered low. After creating virtual markers, implementing virtual marker tracking with saDLC is expected to significantly address the challenges of missing predicted keypoints and ID switches in scenes with occlusion and crowding, enabling high-accuracy tracking. Overall, vmTracking appears to provide a higher level of accuracy with less effort than markerless multi-animal tracking alone, despite requiring modification work to create virtual markers and an additional tracking process.

### Tracking when the number of subjects changes

In maDLC, when the number of target subjects in the video frames changes, keypoints of undetected subjects are not predicted. However, in saDLC, all keypoints are predicted regardless of the number of subjects present. This feature helps to prevent the missed predictions that frequently occur in maDLC during occlusion and crowding, contributing to the enhanced accuracy of vmT-DLC. However, when the actual number of subjects changes, saDLC inevitably results in stray keypoints predicting alternative locations, which can potentially cause confusion in the tracking results.

Additionally, there is a fundamental dilemma in how to manage these stray keypoints during refinement for retraining. To avoid the constant effort of selecting and deleting these inevitably occurring stray keypoints, we attempted to apply them to the remaining mice during refinement. Importantly, this process involves labeling incorrect mice, which have virtual markers different from the correct ones that should be used for labeling the stray keypoints. When no target mouse is present, stray keypoints detect a provisional target, but when the correct mouse appears, it is prioritized for tracking. By annotating multiple-colored virtual markers, vmT-DLC can track different markers based on the situation. Consequently, vmT-DLC can handle stray keypoints without significantly compromising accuracy. Virtual marker colors are crucial for individual identification but are not absolute markers. This flexibility may contribute to vmT-DLC’s ability to achieve high accuracy, even if the virtual markers are not strictly accurate.

This validation involved intentionally controlling the entry and exit of subjects within the video frame. It did not address unpredictable movements, such as when only specific body parts (e.g., the head) temporarily left the frame or were entirely obscured by an object. In this study’s validation, there were also instances of dancers overlapping during tracking, making it challenging to predict the occluded dancer’s body parts. Although the tracking results in these cases might not be significantly incorrect, there is room for debate on their precision. Therefore, in these situations—namely, if the target is frequently obscured by physical barriers or frequently enters and exits the camera frame, or if reducing false detections and focusing solely on accuracy within the visible range is prioritized—choosing vmTracking may not be the optimal choice. However, if the study focuses on interaction behaviors in situations where individuals are in close proximity and there is a need to address occlusion and crowding, vmTracking would be the best choice.

### Applicability to pose analysis tools

With advancements in machine learning-based pose tracking technology, various tools for analyzing pose tracking data obtained from systems such as DeepLabCut, SLEAP, and OpenPose have also evolved, including Keypoint-MoSeq [18], DeepOF [20], B-SOiD [21], and A-SOiD [22]. These machine learning-based tools are powerful for analyzing lateral tracking results and extracting behavioral features, including posture. They vary in approach, with some utilizing unsupervised learning, such as Keypoint-MoSeq, and others using supervised learning to assign behavioral labels, such as A-SoiD, which can also operate in unsupervised mode. However, regardless of the method, if the tracking data contained a high level of noise, then the risk of extracting incorrect behavioral features increased, regardless of how advanced the analysis tool is. For example, evaluating scenes with occlusion or crowding is essential for analyzing interactive behaviors; however, tools such as maDLC and SLEAP are prone to prediction omissions and ID switches in such scenes. To fully leverage these pose analysis tools and achieve reliable behavioral extraction, improving pose tracking accuracy is crucial. The improvement in tracking accuracy provided by vmTracking can contribute to supporting the effective use of these pose analysis tools.

In this study, we conducted an evaluation using Keypoint-MoSeq as an application of pose analysis tools. Consequently, we found considerable agreement between maDLC and vmT-DLC. While the observed agreement might seem relatively low given that the same video was analyzed, the frequency of mismatched combinations in the confusion matrix was generally below 1%. Even in the case of syllable IV and syllable VI, which were among the more frequent mismatched combinations, the similarity dendrogram indicated that these syllables were closely related, suggesting that their action patterns do not clearly represent a discrepancy. Overall, the model generation by Keypoint-MoSeq was barely affected by the results from maDLC or vmT-DLC. The difference in accuracy between maDLC and vmT-DLC may not be a significant issue for Keypoint-MoSeq, which may be able to bridge the differences in tracking accuracy by filtering out keypoint noise and extracting behaviors through unsupervised learning.

However, there were instances where neither data set appeared to fully capture movement patterns, and there were cases where maDLC identified postures that did not align with the actual body (or body axis) orientation. As behavior is extracted from time-series data as a pattern of features, it is not always expected to match the actual behavior in every scene. Yet, if the number of errors increases, discrepancies between the extracted behaviors and actual movements may grow, even if keypoint noise is addressed. Generally, keypoint noise is often detected as high-frequency fluctuations in coordinates; therefore, noise arising solely from FPs is expected to be relatively easy to identify. However, in scenes involving prolonged occlusion or crowding, FN errors are more likely to persist, making it difficult to detect noise based on high-frequency fluctuations. Such scenes, with minimal movement, may impact posture pattern extraction more than movement pattern extraction. While vmT-DLC does not guarantee the absence of incorrect behavioral feature extraction, its characteristic of not producing FN errors may be advantageous for filtering keypoint noise accurately. Consequently, although the effect may be small, vmT-DLC might be able to capture behavioral features more reasonably than maDLC.

### Cross-species and setting applicability of vmTracking

The efficacy of vmTracking across various species and environments has been validated in both fish and humans. Tracking fish schools is considered a challenging task with maDLC [7], often resulting in noisy outputs characterized by frequent keypoint switches, which was observed in this study.

However, implementing vmTracking mitigated these instabilities in maDLC tracking. For human tracking, vmTracking was applied to human dancing, but occasional failures in individual keypoint tracking were still observed during movement, suggesting the need for further improvements, such as refining the placement of virtual markers. Nonetheless, vmT-DLC did not exhibit keypoint set-level ID switches, where entire sets of keypoints switch rather than individual keypoints.

Additionally, in low-movement scenes, tracking failures were rare despite occlusion and crowding. Thus, vmT-DLC demonstrated relatively stable accuracy in tracking the postures of multiple dancers compared to that in maDLC, which often encounters keypoint set-level ID switches and tracking errors even in low-motion scenes. This suggests vmTracking’s potential application in human scenarios such as sports analysis, including highly intermixed contact sports like soccer and basketball, where player interactions are frequent.

In human pose estimation, tools like OpenPose [23] and AlphaPose [24] are highly regarded for human tracking. However, human pose estimation also faces challenges such as occlusion and crowding, with various improved approaches being explored to address these issues [25–27]. Occlusion and crowding can lead to ambiguous or incorrect individual identification. As vmTracking is particularly effective in addressing these issues, it presents a promising solution for the challenges in the human pose estimation field, though there is still room for improvement in the accuracy of individual keypoints.

### Optimizing virtual marker creation

Virtual markers have been shown to serve a role equivalent to physical markers. In addition to not requiring consideration of potential behavioral impacts, as with physical markers, virtual markers also offer the advantage of greater flexibility in their number and placement. This raises the question of how best to configure virtual markers.

We used two virtual markers for mice and fish to ensure redundancy in case one is missing. Setting two virtual markers also helps clarify individual identification in scenarios where two animals overlap. However, tracking results from virtual marker videos created with idtracker.ai suggested that high tracking accuracy can also be achieved with just one virtual marker per animal. Furthermore, experimental results with physical markers similarly indicated that a single marker could be effective. The physical markers were positioned off-center from the torso and, while they could be obscured depending on posture, they still proved effective. For human tracking, we assigned six virtual markers to distinguish independently moving limbs and the head, further clarifying individual identification (Fig 8B). However, for human tracking, using just two points—the head and trunk—achieved comparable accuracy to using six virtual markers. Nevertheless, although the statistical difference was not significant, tracking accuracy generally improved with more virtual markers, suggesting that increasing the number of markers may provide some benefit, even if limited.

While it might seem beneficial to increase the number of virtual markers, doing so may shift them from serving as individual identifiers to acting as indicators of body parts, which could occur even with only a few markers. For instance, if a virtual marker is set to a keypoint corresponding to the head in markerless tracking, most of the virtual markers will generally be located near the head. However, if a marker occasionally appears in a different location, that location might be misidentified as the head. This is an inevitable issue, but it might be that the likelihood of such misidentifications increases as the number of virtual markers increases. As the target is the animal itself, not the virtual marker, it is preferable to use fewer markers to better capture the natural characteristics of the subject.

When creating virtual marker videos, tracking keypoints that will not be used as virtual markers is theoretically unnecessary. For example, in our human tracking experiments, tracking parts like wrists or ankles was unnecessary. Thus, we attempted to create virtual markers with minimal keypoint tracking with maDLC in mouse experiments but could not obtain predictions, as tracklet stitching failed. This suggests that tracking a larger number of keypoints, even if they are not intended as virtual markers, may yield better results for the creation of virtual markers. Additionally, although even a single virtual marker can improve accuracy, tracking a larger number of keypoints in markerless tracking seems beneficial for accommodating various virtual marker configurations. Although having many unnecessary keypoints complicates manual corrections in virtual marker creation, the process can be streamlined by deleting unnecessary keypoints and focusing solely on those intended for virtual markers (S10C and S10D Fig). Alternatively, employing individual tracking tools such as idtracker.ai [19] with single-point virtual markers may allow faster and easier creation of virtual markers than using pose tracking tools.

### Limitations

While vmTracking can be considered a method for high-accuracy tracking of multiple animals, it is important to recognize that there are few limitations that can make the application of vmTracking challenging.

First, vmTracking relies on virtual markers for individual identification, which makes applying vmTracking nearly impossible in situations where there are too many targets to be identified by virtual markers alone. However, such situations are mostly encountered in natural environments with wild animal herds, and vmTracking should present no issues in controlled environments like laboratories.

Second, from the perspective of preparing virtual marker videos, applying vmTracking becomes challenging for videos that are extremely difficult to manage with markerless multi-animal tracking because creating virtual markers becomes difficult. Specifically, it is challenging in cases where many frames fail to detect most targets, as seen in scene V of Fig 7F. However, there is no restriction on which tools to use for creating virtual markers. As different tools may yield better tracking results depending on the video, there are already various markerless multi-animal tracking tools [28–30] in addition to maDLC [7], SLEAP [8], and idtracker.ai [19], potentially offering solutions to overcome this challenge.

Third, when the subject is completely obscured by a large object, this is essentially equivalent to the subject not being present in the video frame, and as with other tools, tracking is impossible. vmTracking is not a method for tracking something that cannot be seen.

Fourth, while vmTracking using saDLC offers the advantage of eliminating FNs, it predicts even in scenarios where it would be preferable not to, such as in unpredictable situations or when the subject is temporarily out of the video frame. Currently, there is no effective method to address this issue, and the only practical solution involves manually removing the corresponding keypoints.

In summary, the development of vmTracking in this study marks an advancement in the field of markerless multi-animal pose tracking. vmTracking has demonstrated exceptional accuracy and efficiency, particularly in addressing challenges such as occlusion and crowding. The reliance on virtual markers simplifies the tracking process, greatly reducing the need for manual annotation and training, which makes vmTracking more user-friendly and practical for laboratory settings and broader applications. The efficiency of vmTracking allows for high accuracy with relatively few annotation frames, highlighting its potential for animal tracking in complex environments. Further research into the impact of the number, color, size, and position of virtual markers on tracking accuracy will open up new possibilities for refining vmTracking. Overall, vmTracking is a robust alternative to traditional tracking methods and a useful tool in the study of animal behavior, ecology, and related fields, providing an effective and efficient solution to some of the most persistent challenges in multi-animal tracking.

## Materials and Methods

### Multi-mouse dataset

Behavioral data were collected from C57BL/6J mice housed in groups of two to three per cage in an environment maintained at a temperature of 24–26 °C and under a 12-h light-dark cycle. These mice had unrestricted access to food and water. All behavioral recordings were performed during the light phase of the cycle. Cages and housing conditions were refreshed weekly. All experimental procedures received approval from the Institutional Animal Care and Use Committee of Doshisha University.

Data collection occurred in two different environments. In the first setting, five C57BL/6J mice freely roamed a rectangular open field with a black floor measuring 55 × 60 cm and walls 40-cm high. Recordings in this environment were captured at a resolution of 640 × 640 pixels, a frame rate of 30 fps, and lasted approximately 30 min, with a total of four videos recorded. In the second environment, three C57BL/6J mice were placed in a circular open field with a white floor measuring 31 cm in diameter and transparent walls 17 cm high. Recordings in this setting were conducted at a resolution of 460 × 460 pixels, a frame rate of 30 fps, and lasted approximately 60 min, with a total of four videos recorded.

Additionally, four videos were recorded where the number of mice in the video frame was manipulated by introducing or removing mice from the arena at intervals of about 12 min. In this session, temporary IDs were assigned to three mice, with the sequence as follows: three mice for the first 12 min, two mice (IDs 1 and 2) for the next 12 min, one mouse (ID 1 only) for the subsequent 12 min, two mice (IDs 1 and 3) for the next 12 min, and finally three mice for the last 12 min, as depicted in S5C Fig. The recording of the physical marker video for the three mice was conducted using the same experimental setup as described for the three mice above, at a resolution of 464 × 464 pixels and a frame rate of 30 fps, for two 30-min sessions. Since the physical markers occasionally detached from the mice during recording, 36 min of data, during which the physical markers remained attached, were extracted for analysis.

### Multi-fish dataset

We used three previously published videos featuring 10 fish swimming in a rectangular aquarium, recorded at a resolution of 2456 × 2058 pixels and 30 fps [31]. For our analysis, the videos were scaled down to 1228 × 1030 pixels. The videos are available at: https://drive.google.com/drive/folders/1Xn3IY46t7DkG2bntbO6sL3oElRPeQA1d.

### Multi-human dance dataset

We used a video from a publicly available dataset featuring seven dancers wearing uniforms, recorded at a resolution of 1280 × 720 pixels and 20 fps [32]. The video is available at: https://drive.google.com/drive/folders/1ASZCFpPEfSOJRktR8qQ_ZoT9nZR0hOea.

### vmTracking procedure and tracking tools

The basic procedure for vmTracking is detailed in the protocol provided in the supplementary information (S1 Protocol and S8–S12 Fig). In this study, we used DLC version 2.2.3, SLEAP version 1.3.3, and idtracker.ai version 5.2.12 for vmTracking.

For basic usage of these tools, please refer to the documentation provided for each tool. For DLC, in the maDLC project, we utilized DLCRnet as the training network because it is known for its high performance in tracking multiple subjects (7), with a maximum of 200,000 training iterations. Conversely, for single-animal DLC (saDLC) used in virtual marker tracking, EfficientNet_b0 was selected as the training network due to its superior performance and lower computational costs compared with ResNet or MobileNet [33]. The maximum number of training iterations was set to 500,000 for the mouse six-keypoint tracking experiment, and to 200,000 for other experiments.

Regarding SLEAP, we applied the top-down method for multi-animal tracking in the mouse six-keypoint tracking experiment and the fish five-keypoint tracking experiment, while both top-down and bottom-up methods were applied for the mouse lateralized keypoint tracking experiment. For single-animal virtual marker tracking in SLEAP, the single method was employed in the analysis pipeline.

### Mouse six-keypoint tracking experiment

For markerless tracking, separate maDLC projects were created for tracking five mice and three mice. Frames were extracted using k-means clustering from videos of five mice (four videos) and three mice (eight videos, including videos where the number of mice in the frame was manipulated). Six points along the midline from the nose to the base of the tail were annotated for each mouse (S1 Protocol and S9 Fig, maDLC labeling). Skeletons were constructed for all keypoint pairs. In this study, annotations were made based on estimations even when body parts were hidden due to occlusion or crowding, across all experiments. These estimations were primarily based on visible body part information (e.g., if the positions of the head and base of the tail were visible, the torso could be estimated); however, when the estimation was challenging using only visible data, temporal movement in the video was also reviewed to support the estimation.

After the initial training, performance evaluation and video analysis were conducted. Based on the results, outlier frames were extracted using the jump algorithm, annotated, and then further trained. In maDLC, each additional training session following refinement was counted as an “iteration” (with the first training before refinement labeled as iteration-0 and the training after the first refinement as iteration-1). In the 5-mice tracking experiment, 7 iterations were performed, resulting in a total of 2,097 annotated frames. In the 3-mice tracking experiment, 11 iterations were performed, resulting in a total of 3,886 annotated frames. Data for tracking accuracy evaluation were obtained from these trained datasets. These annotation data generated by maDLC were also imported into SLEAP and used as training data for tracking.

For reference, the number of frames extracted per iteration in maDLC was set to 50 frames per video. However, there were cases where previously extracted frames were re-selected as outlier frames, and some videos were excluded from extraction (for example, in the three-mouse tracking experiment, we did not extract frames from videos in which the number of mice in the arena was manipulated starting from iteration-7 onward). Therefore, a perfect proportional relationship is not found between the number of iterations and the number of annotated frames. In this study, the number of iterations is used as an intermediate point for comparing results based on the number of annotated frames, and the effect of the iteration count itself is not taken into consideration.

### Creating virtual marker videos

In vmTracking, it is necessary to conduct an independent markerless tracking project to assign virtual markers for individual identification. However, in this study, as there was an existing markerless tracking project for comparison with the vmTracking results, instead of creating a separate project for generating virtual markers, we used the aforementioned markerless tracking project for virtual marker creation. The virtual markers were created based on the intermediate data obtained during the additional training aimed at refining the markerless tracking results. Note that corrections, such as ID switches, were made during the virtual marker creation, but these annotations were independent of the annotations used to improve the multi-animal tracking results. No additional training for markerless tracking was conducted using the corrected results for virtual marker creation.

Virtual markers were created based on the markerless tracking results by correcting ID switches and keypoint position deviations to ensure consistent identification of individuals throughout the video (S1 Protocol and S10A and S10B Fig). The corrected results were then output as a tracking video. The only requirement during this process was to ensure that the tracking remained mostly consistent throughout the video. Therefore, in cases where an ID switch occurred and reverted a few frames later (e.g., within two frames), corrections might not have been made if it was overlooked or if it was judged that the deviation from the correct ID was not significant.

Virtual marker videos for the 5-mice tracking experiment were created based on the SLEAP results with 1,097 annotated frames. From the predicted results of markerless multi-animal pose tracking, all predicted keypoints except for keypoints 2 and 4, which served as virtual markers (S1 Protocol and S11A Fig), were deleted. Subsequently, corrections were made to the ID switches and keypoint positions. Frames in which only one mouse was predicted were extracted, annotations for the four unpredicted mice were added, followed by retraining and re-prediction. Additionally, frames in which only two mice were predicted were extracted. Random frames were selected, annotations for the three unpredicted mice were added, and further training and predictions were performed.

Based on these results, corrections for ID switches and keypoint positions were made as described above to ensure that IDs remained as consistent as possible throughout the video while checking the data using the seek bar at the bottom of the SLEAP GUI. At this stage, labels for the unpredicted mice were not added. Using these corrected data, labeled videos were created as virtual marker videos. In the SLEAP GUI’s view menu, the node size was set to 2, the color palette was set to “standard,” and edges were not displayed.

Virtual marker videos for the three-mice tracking experiment were created based on the results from iteration-6 of maDLC with 2,820 annotated frames. For videos where the number of mice present in the video frames was varied by removing or reintroducing mice into the arena, the virtual marker videos were created based on the videos analyzed with iteration-11 of the training data with 3,886 annotated frames. These videos were corrected using the “Refine Tracklets” function in maDLC, which allows ID switches and keypoints to be manually adjusted (see Protocol S1 for details). Here too, as mentioned above, corrections were made to ensure that consistent IDs were assigned throughout the video by adjusting ID switches and keypoint positions. The sections requiring correction due to ID switches or keypoint deviations were identified by reviewing the pre-correction tracking video. Using these corrected data, labeled videos were created as virtual marker videos. In scenarios where the number of mice in the video frame changed, overlapping tracking occurred due to a reduction in the number of individuals. To address this, a custom GUI (S10C Fig) was used to delete the coordinate data of IDs not used as virtual markers before creating the video. The keypoints in these videos were displayed using the matplotlib colormap “gray,” with a marker size set to 2 (as specified in the configuration file). Only keypoints 2 and 4 of each mouse were used as virtual markers (S1 Protocol and S11A Fig), and no skeletons were outputted.

Furthermore, to simplify the creation of virtual marker videos, we also conducted an independent maDLC project to track only the two keypoints intended to be designated as virtual markers. However, during the initial video analysis step, tracklet stitching failed in all videos, and we were unable to obtain coordinate data, leading to the project’s discontinuation at that stage.

### Virtual marker tracking with single-animal tracking methods

A saDLC project was created for virtual marker tracking with these virtual marker videos. However, as saDLC is not designed for multi-animal tracking, the number of individuals and tracking points cannot be set to reflect multiple animals. Therefore, the total number of keypoints was set by multiplying the 6 keypoints per mouse by the number of tracked mice (i.e., a total of 30 keypoints for 5 mice and 18 keypoints for three mice). Refer to Appendices S1 and S3 for the differences in settings between maDLC and vmTracking (saDLC), particularly regarding individuals and bodyparts, and refer to Appendices S2 and S4 for the differences in the header of the coordinate data files resulting from the different projects. Each mouse was allocated six keypoints (along the midline from the nose to the base of the tail), and skeletons were constructed for all keypoint pairs. For annotation, 50 frames were extracted by k-means clustering from each video, and annotations were made by assigning consistent IDs across frames, based on the color of the virtual markers (S1 Protocol and S9 Fig, saDLC labeling), followed by training and analysis. In the 5-mice tracking experiment, 5 iterations were performed, resulting in a total of 1,196 annotated frames. In the 3-mice tracking experiment, 1,167 frames were annotated across 5 iterations using 4 videos with a fixed number of mice in each frame, totaling 5 iterations. Data for tracking accuracy evaluation were obtained from these trained datasets. These annotation data generated by saDLC were also imported into SLEAP and used as training data for virtual marker tracking.

For the analysis of scenarios where the number of mice in the video frame changes, frames from these scenarios were added to the initial 1,167 annotation frames for additional training. In saDLC, the keypoints specified in the configuration file are almost always predicted without exception, so that no missing keypoints occur. Consequently, when the number of tracked individuals in the arena decreases, stray keypoints appeared as they lost their tracking targets. To deliberately manage these stray keypoints, the remaining mice were annotated with stray keypoints to learn new tracking rules. Specifically, in frames where only two mice were present in the arena, labeling was performed such that IDs 2 and 3 overlapped. However, in frames where only one mouse was present, all labels were assigned to that single mouse, an additional 517 frames were added (for a total of 1,684 frames), and data for tracking accuracy evaluation were obtained from these trained datasets.

### Mouse lateralized keypoint tracking experiment

In the lateralized keypoint tracking experiment, we conducted both a physical marker tracking experiment and a vmTracking experiment. To eliminate the influence of factors such as differences in the annotation data used for training across conditions, the number of annotated frames, and the number of iterations, the same annotation dataset was prepared. Additionally, no refinement or additional training was performed, and the results after a single round of training were compared. When using DLC, although there was a difference in the training network—DLCRnet for maDLC and EfficientNet_b0 for saDLC—the maximum number of training iterations was unified at 200,000.

In the physical marker videos, there were instances where the physical markers detached from the mice. From the sections of the video where the markers remained attached, 12 3-min video clips (i.e., each consisting of 5,400 frames) were extracted. Using the physical markers as a guide, we annotated 14 keypoints per mouse on 180 frames per clip, totaling 2,160 frames, and trained the data using the same annotation dataset with maDLC and saDLC. In the physical marker tracking experiment, tracking results were obtained by analyzing the video clips used for training.

In the vmTracking experiment, we used the markerless video and corresponding virtual marker video from the three-mice, six-keypoint tracking experiment. Additionally, we tracked the markerless video using idtracker.ai (described later) and created another virtual marker video (ID virtual marker video) that was distinct from the one created with maDLC. From each of these three types of videos, 24 3-min video clips (i.e., each consisting of 5,400 frames) were extracted from the same scenes.

Using the virtual marker videos, we annotated 180 frames per video clip, totaling 4,320 frames. This annotation dataset was then divided into two independent datasets, each containing 12 clips. The two datasets were trained independently, with the markerless video trained using maDLC, and the virtual marker and ID virtual marker videos trained using saDLC. After training, tracking results were obtained by analyzing the video clips from the dataset that was not used in training, thereby minimizing the effects of overfitting on the training data. Additionally, the annotation data were imported into SLEAP, where the markerless video was trained using the top-down and bottom-up methods, and the virtual marker and ID virtual marker videos were trained using the single method. As with DLC, videos that were different from the training dataset were analyzed. The results were used as the data for tracking accuracy evaluation.

### Fish tracking experiment

A maDLC project was created for tracking 10 fish, with 50 frames extracted from each of 3 videos using k-means clustering and 5 evenly spaced points annotated along the midline from the head to the tail tip. These data were imported into SLEAP for training and analysis, and corrections were made based on these data to create a virtual marker video. The second and fourth keypoints from the head side were used as virtual markers, with node size set to 2 and the color palette set to “alphabet.”

A saDLC project was created for virtual marker tracking with these virtual marker videos. Five keypoints per fish were set, totaling fifty keypoints assigned across ten fish. Annotation was performed with consistent IDs assigned across frames based on virtual marker color, followed by training, analysis, and refinement, resulting in a total of 852 annotated frames (no tracking accuracy evaluation data were obtained at this stage). Next, to compare maDLC and vmTracking under similar conditions, this annotation dataset was applied to each project, followed by training and analysis to obtain results for tracking accuracy evaluation. In this study, a single video consisting of 2,259 frames (with 410 annotated frames) was used for evaluation.

### Dance tracking experiment

A maDLC project was created to track 7 dancers, with 20 frames extracted using k-means clustering and 19 keypoints annotated, including the head, neck, trunk, shoulders, elbows, wrists, hands, hips, knees, ankles, and feet. Based on the results obtained from 59 frames after repeated training, analysis, and refinement annotations, corrections were made to create virtual marker videos. Two types of virtual marker videos were created: one using two points (head and trunk) and another using six points (the same two points plus one point on each elbow and knee) (S1 Protocol and S11B Fig). The virtual marker size was set to 5 and the color map was set to “rainbow” in matplotlib.

A saDLC project was created for virtual marker tracking with the six-point virtual marker video. Each dancer was assigned 19 keypoints, totaling 133 keypoints across 7 dancers. Annotation was performed with consistent IDs assigned across frames based on virtual marker color, followed by training, analysis, and refinement, resulting in a total of 150 annotated frames (as with the fish tracking, no tracking accuracy evaluation data were obtained at this stage). Then, to compare maDLC and vmTracking for the two types of virtual marker videos under similar conditions, this annotation dataset was applied to each project, followed by training and analysis to obtain results for tracking accuracy evaluation.

### Creating tracking videos excluding virtual markers

Tracking videos typically display tracking results as labels on the video used for tracking. In the case of vmTracking, this results in a video where labels are overlaid on the virtual marker video.

However, in this study, we displayed the tracking results on the original markerless video, producing tracking videos without virtual markers (S1–S8 Video). To this end, we replaced the virtual marker video in the tracking project with the original video before generating the final tracking video (S1 Protocol and S12 Fig).

### idtracker.ai

Two markerless videos used in the three-mice tracking experiment were tracked using idtracker.ai. To eliminate the influence of black areas in the background, a region of interest was set to match the field in the markerless videos. The parameters for tracking were set to 3 for Number of animals, with Dark animals selected for a Bold intensity threshold of 100 and Bold area threshold of 1000. After correcting ID switches in the tracking results, the data were output as virtual marker videos, from which 24 3-min video clips were extracted. For tracking accuracy evaluation using centroids, the uncorrected data were used.

### Physical marker

The base apparatus for the physical marker was designed using Onshape (https://www.onshape.com/en/), a cloud-based 3D CAD service, and fabricated with a 3D printer (Form3, Formlabs Inc). Circular colored tack stickers were then applied to the apparatus to serve as physical markers.

### Mice tracking performance evaluation

The accuracy of multi-mouse tracking with six keypoints was evaluated based on a manually created ground truth for sampled scenes, where each scene consisted of either 150 (5 s) or 300 (10 s) consecutive frames. Consecutive frames were used to allow for the evaluation of ID switches. In the 5-mice tracking experiment, 12 OC scenes were evaluated, while in the 3-mice tracking experiment, 12 OC scenes and 8 nOC scenes were assessed. In this study, OC scenes were defined as instances where overlapping occurred in the body regions of two or more mice, while nOC scenes were defined as instances where no such overlapping occurred. The sampled scenes were chosen by reviewing the videos and selecting dynamic scenes whenever possible. In the five-mice tracking experiment, three scenes were selected from each of four videos. In the three-mice tracking experiment, five scenes were selected from each of four videos, which had not been manipulated in terms of the number of mice present in the frame, and these scenes were classified into OC and nOC scenes. This evaluation focused on the accuracy of each keypoint and the centroids derived from them. To compare predicted data with ground truth, each predicted mouse ID was paired with the most matching ground truth ID. For this purpose, distances for all combinations of predicted data and ground truth IDs were calculated, and the Hungarian algorithm was used to determine the most suitable combinations.

### Keypoint evaluation

The predicted points were defined and classified based on whether they were detected within a predefined threshold (10 pixels) from the corresponding GT keypoints (S1 Fig). If a predicted point was within the threshold of the corresponding GT keypoint, it was classified as a “Match”; if it was outside that threshold, it was classified as a FP; and if there was no predicted point, it was classified as a FN. FPs were further classified as follows: those outside the threshold of all GT keypoint (Deviating FP), those matching the predicted body part but with an incorrect ID (ID mismatch FP), and those not matching the predicted body part regardless of ID (Bodypart mismatch FP). When a predicted point fell within the threshold of any GT keypoint (i.e., classified as a Match, ID mismatch FP, or Bodypart mismatch FP), it was marked as an ID switch if its ID had changed from the previous frame.

These metrics were calculated as percentages relative to the number of target points (equivalent to the GT count) and recorded as Tgt match, FN, each type of FP, and ID switches. Additionally, as accuracy metrics focused on the predicted data, we provided Pred match (percentage of Match counts relative to the number of predicted data points) and Bodypart match (the percentage of the sum of Match counts and ID mismatch FP counts relative to the number of predicted data points). RMSE, which is a metric based on the number of predicted data points, was also calculated based on the distance to the corresponding GT keypoint.

In this study, Tgt match was given the highest priority. However, when an ID switch occurred, even if accurate tracking was performed with the switched ID afterward, it may still be classified as a body part mismatch FP, potentially leading to underestimated tracking accuracy. Therefore, to capture matches in cases of accurate tracking after an ID switch, we included a body part match, which was defined as a match that does not require correct ID assignment. While it would have been possible to evaluate the results after making manual corrections, this evaluation method was chosen to more accurately assess the raw outcomes of multi-animal tracking and vmTracking. Notably, because body part match is based on the predicted keypoints, it serves as a more lenient matching criterion compared to the stricter Tgt match. This approach aims to provide accuracy assessments across various criteria.

### Centroid evaluation

For centroid evaluation, the centroid derived from posture-tracking keypoints and the centroid obtained from idtracker.ai were compared based on GT data. In the 14-keypoint lateralized tracking experiment, the centroid was calculated using 11 predicted keypoints, excluding the tail. If only one predicted keypoint was available, then that single point was used as the centroid. A direct comparison with the GT centroid was not performed. This decision was made because the centroid derived from posture tracking keypoints serves as a representative value of the posture tracking data and cannot be said to have the same meaning as the idtracker.ai centroid, making a direct comparison inappropriate. Instead, the obtained centroid was compared with the three central points along the GT keypoints’ midline to assess whether it approximately matched the correct mouse.

Specifically, if the obtained centroid was within 20 pixels of any of these 3 GT keypoints, it was classified as a Tgt match. Considering that the centroid derived from posture tracking and the centroid obtained from idtracker.ai are not equivalent, a larger value was set for the threshold compared to the 10-pixel threshold used in keypoint evaluation. Although the centroids were not directly compared with the GT centroid, as the GT serves as the reference, it was reasoned that the centroid derived from multiple keypoints of posture tracking might be closer to the GT keypoints, thus warranting a larger threshold. If it exceeded this threshold, it was classified as a FP, and if no centroid was obtained (indicating that all keypoints used for centroid calculation were FNs), it was classified as a FN. FPs were further categorized as Deviating FPs and ID mismatch FPs. Unlike keypoint evaluations, body part mismatches could not be assessed; therefore, Bodypart mismatch FPs were excluded from the centroid evaluation. Similarly, only matches based on predicted keypoints (Pred match) were counted, with Bodypart matches excluded. Additionally, since consecutive frame GT data were not used in the lateralized keypoint tracking experiment, ID switches were not counted. RMSE was also excluded from the evaluation, as establishing a common GT centroid was challenging.

### Accuracy of virtual markers

In the mouse tracking experiment, two keypoints in markerless tracking were designated as virtual markers. ID matching for virtual markers was defined as follows: if keypoints 2 and 4, designated as virtual markers, were detected within a threshold distance of 10 pixels from any of the 6 ground truth keypoints of the same mouse, they were considered to have matching IDs (S2A Fig). Based on this criterion, virtual marker assignment patterns were categorized as follows: when both virtual markers were present, both showed matched IDs (Match (Two VM)), only one showed a matched ID (Partial match (Two VM)), or neither showed matched IDs (No match (Two VM)); when only one virtual marker was present (the other missing), then a single ID matched (Match (One VM)) or did not match (No match (One VM)); and when neither virtual marker was present (Missing (No VM)) (S2B Fig).

### Keypoint-MoSeq

To assess whether differences in accuracy between maDLC and vmT-DLC affect the results of pose analysis tools, we applied data from these methods to Keypoint-MoSeq, which is a machine learning tool that automatically identifies behavioral modules (syllables) from keypoint data and filters out noise of keypoints to enhance syllable identification. Nonetheless differences in tracking accuracy may still influence syllable identification.

We used results from a three-mice tracking experiment with six keypoints for evaluation using Keypoint-MoSeq. First, ID switches were corrected (excluding instances where IDs quickly reverted) to ensure ID consistency within the dataset because ID switches could have a significant impact on Keypoint-MoSeq’s module identification. Then, a model was created using maDLC and vmT-DLC results derived from one of these videos. For model training, we ran 50 iterations of AR-HMM fitting and 500 iterations of full model fitting with vmT-DLC data, followed by 200 iterations with maDLC data. The κ value for full model fitting was set to 1e4 (10,000). This model was then applied to data from both maDLC and vmT-DLC, and results were assessed. Only syllables occurring at a frequency of 1% or higher were output, covering a movement duration of 1 s in the plot (0.5 s before and after each syllable).

Based on the resulting syllable plots and similarity dendrogram, we grouped three low-movement syllables into a single category. Additionally, syllables with an occurrence rate below 1% (excluded from plots and the dendrogram) were grouped together. Using a confusion matrix of syllables from maDLC and vmT-DLC data, syllable frequencies from both tracking methods were visualized, and Cohen’s kappa coefficient was calculated as an agreement measure. For mismatched syllable combinations with frequencies exceeding 1% in the confusion matrix, cases that remained mismatched for 2 s (60 frames) or more (as brief mismatches might be noise-related) were visually checked against the actual tracking footage to verify validity.

### Evaluation of stray keypoints

In experiments where mice were introduced or removed from the arena, we calculated the distances between identical keypoints for each ID to verify that we could track the remaining mice as temporary targets using stray keypoints that had lost their original targets. Keypoints were considered to overlap if the distance between them was within 10 pixels, and the frequency of such frames was assessed. As overlapping stray keypoints were unnecessary in the cleaned and finalized dataset, we created a GUI-enabled code (https://doi.org/10.5281/zenodo.14249295) to remove these redundant keypoints (S10C Fig). This cleaned and finalized dataset was then used to create the videos.

### Fish tracking evaluation

Of the three videos, a single video consisting of 2,259 frames was used for the evaluation. The evaluation was conducted based on frame-to-frame variations and outliers observed throughout a series of videos. Based on setting equidistant keypoints along the body axis, it was assumed that if the distances between adjacent keypoints were not uniform, the tracking was not accurate. To this end, the variance of distances between adjacent keypoints was calculated as a measure of tracking accuracy, with greater variance interpreted as lower accuracy. The variance for each fish per frame was calculated, the median variance identified, and the frequency of outliers assessed. To consistently assess the frequency of outliers, outlier thresholds were determined using the interquartile range method based on integrated data from maDLC and virtual marker tracking. For each condition, these thresholds were applied, and the number of frames with variance considered outliers for each fish was counted.

Schooling behavior, in which fish often swim in the same direction, was also evaluated. An angle based on 2 anterior keypoints for each fish was calculated to represent the head orientation, and the cosine similarity for head direction among all 10 fish was computed for each frame to examine synchrony. Furthermore, it is rare for the direction of the head to change abruptly between frames, such changes likely indicate a tracking error. Therefore, the angle of head direction was calculated using the two anterior keypoints for each fish, with a focus on the synchronization of these predictions. The difference in angles between frames was measured, and the changes exceeding 90° were counted as abnormal directional changes. Similar frames were interpreted as lower tracking accuracy.

As a sampling-based evaluation using annotation data for training as the GT, the proportions of Tgt matches, FNs, each type of FP, Pred matches, and Bodypart matches were calculated. Additionally, a correlation analysis of cosine similarity was performed based on the number of frames where data were available in maDLC (i.e., frames without missing cosine similarity data).

### Dance tracking evaluation

In this study, we evaluated the detection of similar poses during 2 s of similar movements and 2 s of static poses among seven dancers. For each dancer, we calculated distances between all keypoints to create a standardized distance matrix. The RMSD of the standardized distance matrices between dancers was then computed as a measure of similarity; the more similar the poses, the smaller the RMSD. To determine the similarity of each frame, we calculated the mean RMSD across all dancer pairs per frame. Furthermore, as a sampling-based evaluation using annotation data for training as the GT (150 frames), the proportions of Tgt match, FNs, each type of FP, Pred match, and Bodypart match were calculated.

### Statistical analysis

Outliers in tracking data, often due to tracking errors, are unavoidable. To mitigate the influence of these outliers, all statistical analyses in this study used nonparametric tests. For comparisons between two paired conditions, the Wilcoxon signed-rank test was used. For comparisons among three or more conditions, the Friedman test was applied for paired data and the Kruskal–Wallis test was used for unpaired data, with both followed by Bonferroni correction. Spearman’s rank correlation was used for correlation assessments.

More specifically, metric-by-metric comparisons in the mouse tracking experiments used the Wilcoxon signed-rank test for physical marker data, while other comparisons employed the Friedman test with Bonferroni correction. To examine the relationship between virtual marker accuracy and Tgt match, Spearman’s rank correlation was used. In the OC scene of the three-mice tracking experiment, the accuracy across varying annotation frame counts was evaluated using the Friedman test with Bonferroni correction. Comparisons of accuracy based on annotation frame counts relative to the vmT-DLC level were performed with paired comparisons using the Wilcoxon signed-rank test, applying Bonferroni correction with a comparison count of 12.

For overlapping keypoint evaluations as the number of tracked mice varied, comparisons of d1-3 were conducted within each condition based on differences in the number of tracked mice using the Friedman test (i.e., five comparisons across three conditions). Additionally, comparisons across tracking conditions for each d1-3 were made using the Kruskal-Wallis test (i.e., three comparisons across five conditions), with Bonferroni correction applied to the total comparison count of 45. Due to the small sample size, we also calculated Cliff’s delta as an effect size to aid in interpretation. For the Keypoint-MoSeq analysis, Cohen’s kappa coefficient was calculated to assess syllable agreement between maDLC and vmT-DLC results.

In the fish tracking experiments, the Wilcoxon signed-rank test assessed the median variance in keypoint distances for each fish, the frequency of this variance as an outlier, and the frequency of head orientation changes greater than 90° between frames. Spearman’s rank correlation analysis was conducted to assess the relationship between cosine similarity and ground truth. In the dance tracking experiment, RMSD values based on standardized distance matrices among seven dancers were evaluated using the Friedman test with Bonferroni correction.

The MATLAB functions used for these analyses included friedman (Friedman test), kruskalwallis (Kruskal–Wallis test), multcompare (multiple comparisons), signrank (Wilcoxon signed-rank test), and corr (correlation analysis). Any calculations for which built-in functions were unavailable were performed manually. Statistical significance was set at p < 0.05.

Note that bar graphs were used in the figures when n ≤ 20. In these figures, each plot represented individual measurements, bars indicated the mean, and error bars (if present) indicated the 95% confidence interval. Box plots were used when n > 20. In these plots, the bottom and top of the box represent the first quartile (Q1) and third quartile (Q3), with the line inside indicating the median. Whiskers extended to 1.5 times the interquartile range (Q3 to Q1), and measurements beyond this range are plotted as outliers in circles. Additionally, the mean value is marked with a star symbol.

## Supporting information

S1 Protocol

## Acknowledgements Funding

This work was supported by Kakenhi grants from the Japan Society for the Promotion of Science (21H04247 and 24K15711 to H.A., and 21H05296 and 23H00502 to S.T.) and by the Core Research for Evolutional Science and Technology (CREST) program of the Japan Science and Technology Agency (JPMJCR23P2 to S.T.).

## Author contributions

H. A. and S. T. conceived the study. H. A. acquired the behavioral data of the mice. H. A. performed the animal tracking. H. A. performed the formal analyses. H. A. and S. T. prepared the manuscript.

## Data and code availability

The results of this paper can be reproduced using downloadable data and accompanying code, which are available from the Zenodo repository (https://doi.org/10.5281/zenodo.14249295).

## Declaration of generative AI and AI-assisted technologies in the writing process

During the preparation of this study, the authors used ChatGPT to improve readability and language. After using this service, the authors reviewed and edited the content as needed and take full responsibility for the content of the publication.

## Supplementary information

### Supplementary Figure Legends

**S1 Fig.**
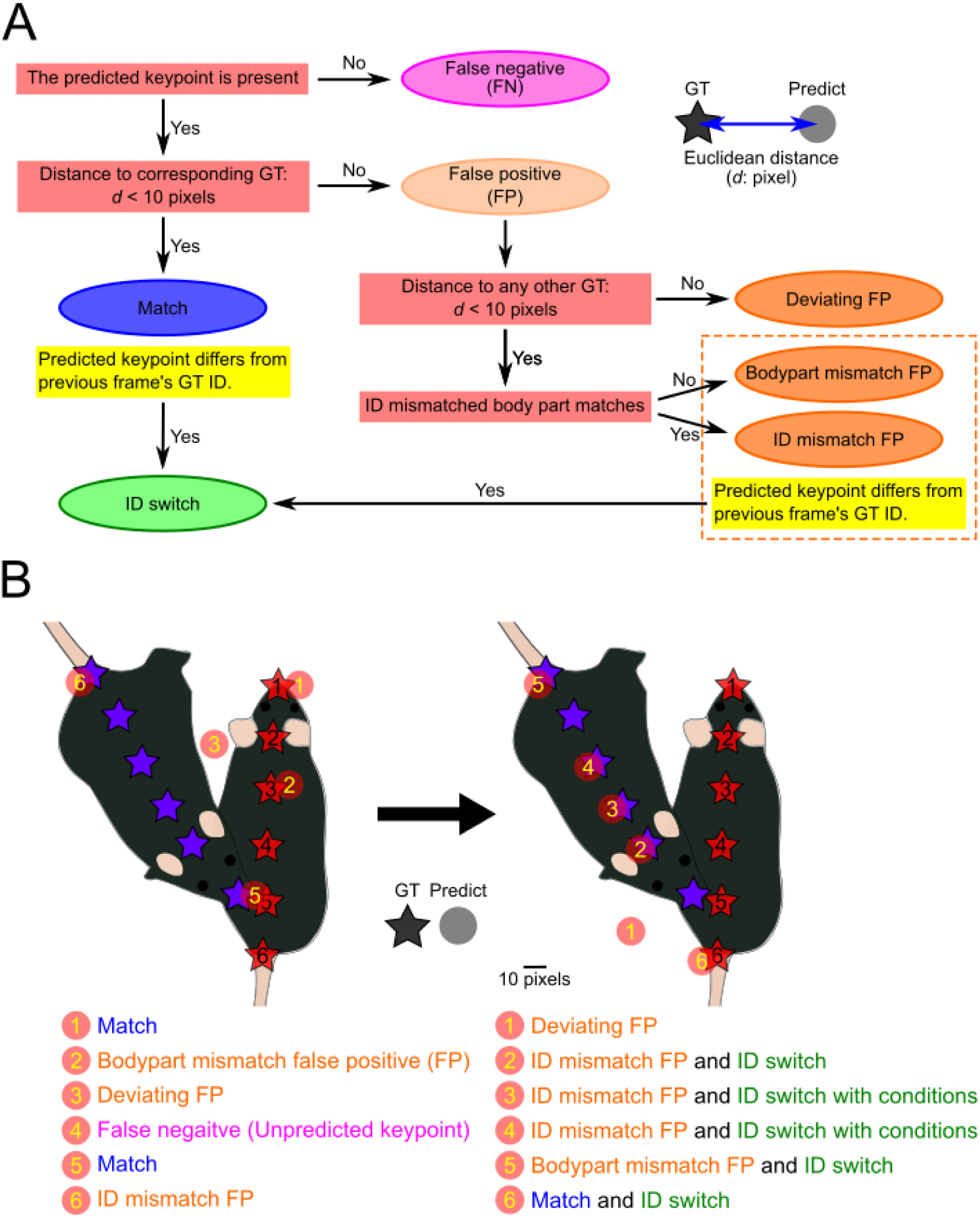
Classification of predicted keypoints for tracking performance evaluation. (A) Workflow of predicted keypoint classification. If a predicted point was within the threshold of 10 pixels from the corresponding ground truth (GT), it was considered a match; if a prediction was made but fell outside this threshold, it was counted as a false positive (FP); and if no predicted point was detected, it was classified as a false negative (FN). Furthermore, FPs were categorized by pattern as follows: FP when outside the threshold for all GT keypoints (Deviating FP); ID mismatch FPs when the body part was correct but assigned to an incorrect ID (ID mismatch FP); and body part mismatch FP when the predicted body part was incorrect, regardless of ID (Bodypart mismatch FP). When a predicted point was within the threshold of multiple GT keypoints, the GT keypoint classified as a match took priority; whereas when no match was identified, the nearest GT keypoint was used for evaluation. If any GT keypoint was within the threshold—such as in the case of a match, ID mismatch FP, or body part mismatch FP—but the ID differed from the previous frame (or an earlier frame if it was a FN and deviating FP), this was classified as an ID switch. Red stars (ground truth) and red circles (predictions) correspond to each other. (B) Example of predictive keypoint classification. For the left of the arrow, 1: accurately predicts a location within 10 pixels of the corresponding GT (Match); 2: predicts a location more than 10 pixels away from the corresponding GT and within 10 pixels of a non-corresponding GT (Bodypart mismatch FP); 3: does not predict within 10 pixels of all GT (Deviating FP); 4: shows no predicted keypoints (FN); 5: is within 10 pixels of both the corresponding and non-corresponding GTs, with the corresponding GT being prioritized (Match); and 6: predicts a location more than 10 pixels away from the corresponding GT and within 10 pixels of a GT with a different ID but matching body part (ID mismatch FP). However, as ID switches are based on changes from the previous frame, it does not qualify as an here. For the right of the arrow, 1: fails to predict within 10 pixels of all GT (Deviating FP); 2: predicts a location more than 10 pixels from the corresponding GT and within 10 pixels of a GT with a different ID but matching body part (ID mismatch FP), and it also predicts a GT with a different ID from the previous frame (ID switch); 3 and 4: predict locations more than 10 pixels from the corresponding GT and within 10 pixels of a GT with a different ID but matching body part (ID mismatch FP). Although these predict GTs with different IDs, in the previous frame, 3 is classified as a deviating FP and 4 as an FN, and since the predicted IDs for the previous frame cannot be determined, the judgment of whether an ID switch has occurred depends on the results from even earlier frames. If the predicted IDs had changed from even earlier frames, then this situation would be classified as an ID switch. 5: Predicts a location more than 10 pixels away from the corresponding GT and within 10 pixels of a GT where both ID and body part are incorrect (Bodypart mismatch FN), and it also predicts a GT with a different ID from the previous frame (ID switch); and 6: accurately predicts within 10 pixels of the corresponding GT (Match), and it also predicts a GT with a different ID from the previous frame (ID switch).

**S2 Fig.**
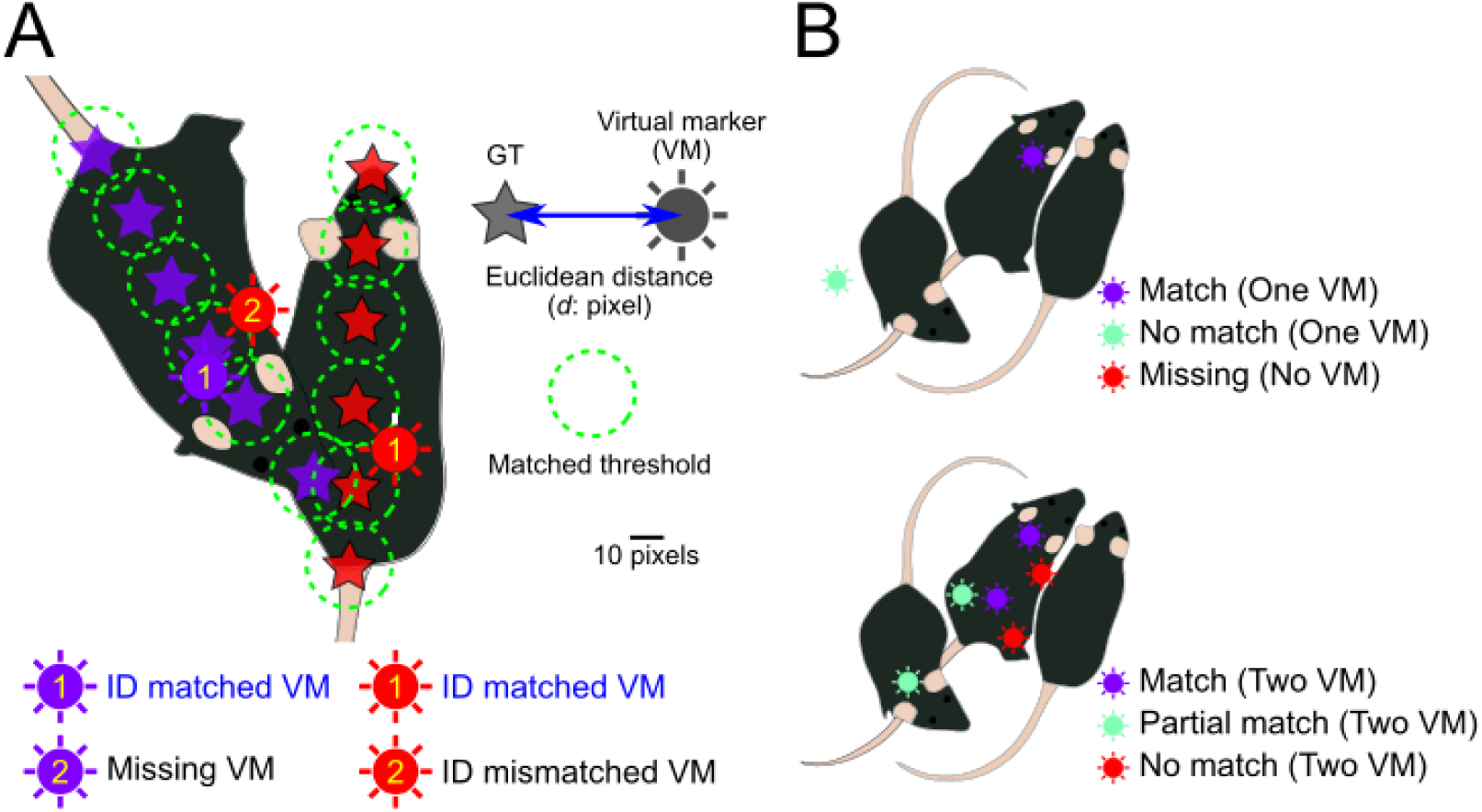
Accuracy of virtual markers. (A) Concept of ID matched virtual markers, which indicates whether a virtual marker’s ID match was based on the distance (*d*: pixels) between the virtual marker points indicated by radial circles and the ground truth indicated by stars. If any ground truth keypoints with the same ID as the virtual marker were within 10 pixels of the corresponding virtual marker, the virtual marker was considered to have an ID match (as shown in the figure for the red and purple virtual markers labeled “1”). Conversely, if no ground truth keypoints with the same ID as the virtual marker were within 10 pixels, the virtual marker was considered to have an ID mismatch (as shown in the figure for the absent purple virtual marker labeled “2”). Additionally, if virtual markers were absent due to missing predictions in markerless multi-animal pose tracking, these were treated as instances of ID mismatch, as they represent missing virtual markers that should have been present (as shown in the figure for the purple virtual marker labeled “2”). The proportion of ID-match virtual marker points for each scene was calculated and used for the correlation analysis with the accuracy of vmTracking. (B) Classification of virtual marker assignment patterns In the mouse tracking experiment, two keypoints in markerless multi-animal tracking were designated as virtual markers. For frames where both keypoints are present, virtual marker assignments are categorized as follows: both IDs match (Match (Two VM)), only one ID matches (Partial match (Two VM)), and neither ID matches (No match (Two VM)). When only one keypoint is present (the other being missing), it is categorized based on whether its ID matches (Match (One VM)) or does not match (No match (One VM)), and when both keypoints are missing, it is classified as Missing (No VM). This classification was performed for each mouse in each frame, and the frequency of each category was calculated for each scene.

**S3 Fig.**
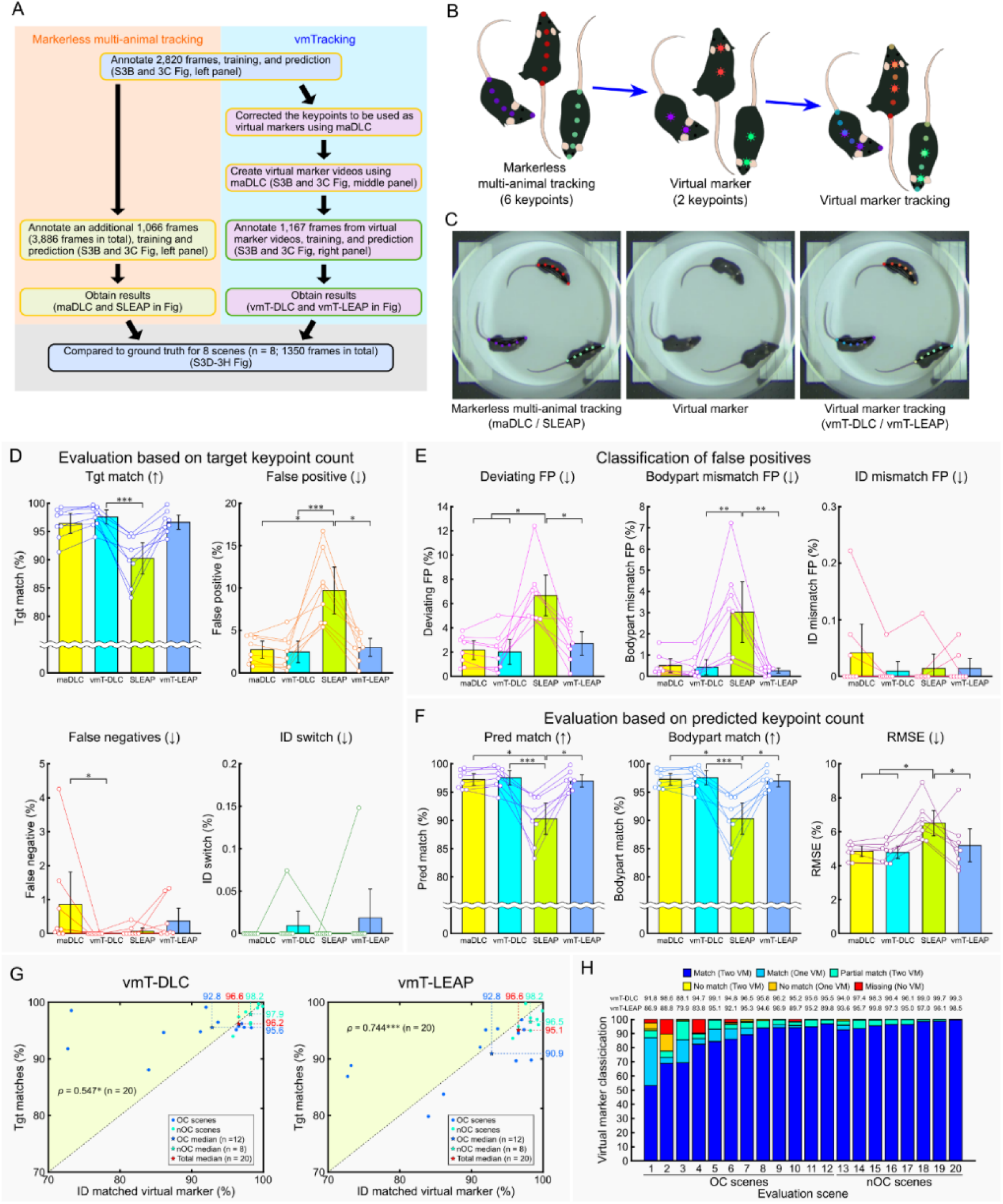
Comparison of markerless multi-animal pose tracking and vmTracking in a non-occlusion-crowded scene of three mice (Related Fig 3). (A) Overview of the verification process. The yellow box in the schematic represents the processes using multi-animal tracking tools, while the green box represents the processes using single-animal tracking tools. (B) Schematic of virtual marker creation. Six keypoints per mouse were tracked, with two designated as virtual markers. (C) Examples of markerless multi-animal pose tracking (left), virtual marker (middle), and virtual marker tracking (right) in an occlusion-crowded scene. (D–F) Various percentage-based metrics were compared across multi-animal DeepLabCut (maDLC), virtual marker tracking with DeepLabCut (vmT-DLC), Social LEAP (SLEAP), and virtual marker tracking with SLEAP (vmT-LEAP), based on manually generated ground truth (GT) data. (D) The evaluation was conducted as a percentage of the target keypoints, with the total number of GT keypoints as the denominator. The percentages of matches (Tgt match), false negatives, false positives, and ID switches were calculated based on the number of GT keypoints. (E) False positives from (D) were classified into deviations exceeding the threshold from all GT (Deviating FP), mismatches in predicted body parts (Bodypart mismatch FP), and cases where the predicted body part was correct but the ID was incorrect (ID mismatch FP). (F) Evaluation was conducted as a percentage of the predicted keypoints, excluding FNs, using the total predicted keypoints as the denominator, and the metrics included matches (Pred match), body part matches (Bodypart match), irrespective of ID, and root mean square error (RMSE). Arrows indicate whether higher or lower values are better. Data are shown as the mean ± 95% confidence interval (n = 8 scenes). Statistical analysis was performed using the Friedman test with Bonferroni correction. (G) Scatter plots illustrating the relationship between ID matched virtual markers and Tgt match for vmT-DLC (left) and vmT-LEAP (right). The plots are color-coded to differentiate OC scenes (light blue) and nOC scenes (light green). The plots in the upper-left (light yellow-green) area indicate cases where Tgt match exceeded the ID matched virtual markers. Filled circles represent individual scenes, and the stars signify their median. Correlation analysis was conducted by combining OC and nOC scenes using Spearman’s rank correlation (n = 20 scenes). All statistical significance levels were set at p < 0.05. *:p < 0.05, **: p < 0.01, ***:p < 0.001. (H) Stacked bar graph showing the proportion of virtual marker assignment patterns in each scene. Virtual markers were classified as follows: when two virtual markers were assigned to a single mouse, both IDs matched (Match (Two VM)); one virtual marker was missing and the other’s ID matched (Match (One VM)); one virtual marker’s ID matched while the other’s ID did not match (Partial match (Two VM)); neither ID matched (No match (Two VM)); one virtual marker was missing and the other’s ID did not match (No match (One VM)); and both virtual markers were missing (Missing (Two VM)). The proportions of each category are displayed by scene. The values for vmT-DLC and vmT-LEAP above the figure represent the Tgt match values for each scene.

**S4 Fig.**
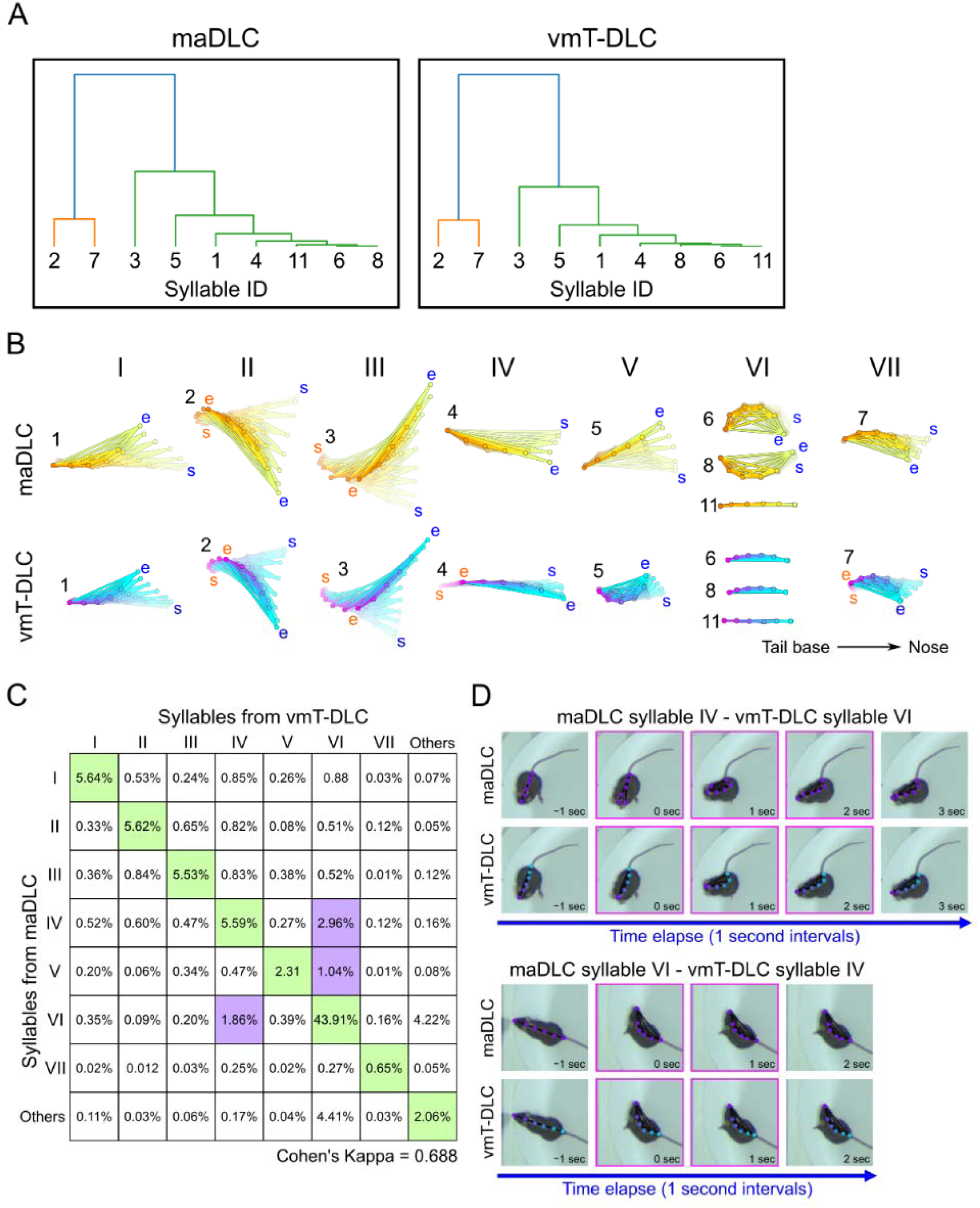
Comparison of keypoint-MoSeq results between multi-animal DeepLabCut and vmTracking. (A) Similarity dendrogram of syllables obtained from keypoint-MoSeq analysis on the tracking results of multi-animal DeepLabCut (maDLC) and vitual marker tracking with single-animal DeepLabCut (vmT-DLC) across all 112,991 frames. The syllable IDs correspond to the Arabic numerals in (B). (B) Behavioral syllables obtained from maDLC and vmT-DLC. Here, syllables 6, 8, and 11 were grouped and processed as nearly immobile patterns based on the similarity dendrogram results and syllable patterns. Each syllable represents a 1-s behavioral pattern, with the left side of each plot representing the base of the tail and the right side representing the head (snout). In the figure, “s” denotes the start point of the behavior, and “e” denotes the end point. Blue text indicates head-side movement, while orange indicates tail-base movement. When there was minimal change at the start or end points, these letters were omitted. (C) Confusion matrix for syllable similarity obtained from maDLC and vmT-DLC. The Roman numerals correspond to the seven syllables in (B), and minor syllables not included in I–VII are grouped as “Others.” The matrix shows the frequency of syllable number combinations assigned to each frame by maDLC and vmT-DLC (for example, frames where syllable I was assigned by both maDLC and vmT-DLC make up 5.64% of the total). Cells shaded in light green indicate frames where the same syllable was assigned by both maDLC and vmT-DLC, while cells shaded in light purple indicate cases where different syllables were assigned by maDLC and vmT-DLC, with a frequency of 1% or more (excluding cases that involve “Others”). The value below the matrix represents Cohen’s kappa coefficient as an indicator of agreement. (D) Examples of tracking by maDLC and vmT-DLC in scenes where the assigned syllables did not match and this discrepancy persisted for more than 2 s (60 frames). The upper row shows a scene where maDLC consistently assigned syllable VI while vmT-DLC assigned syllable IV. The lower row shows a scene where maDLC assigned syllable IV and vmT-DLC assigned syllable VI. Snapshots are shown at 1-s intervals, with the scenes where the syllables were assigned highlighted in magenta, and the frames at both ends corresponding to 1 s before and after the scene.

**S5 Fig.**
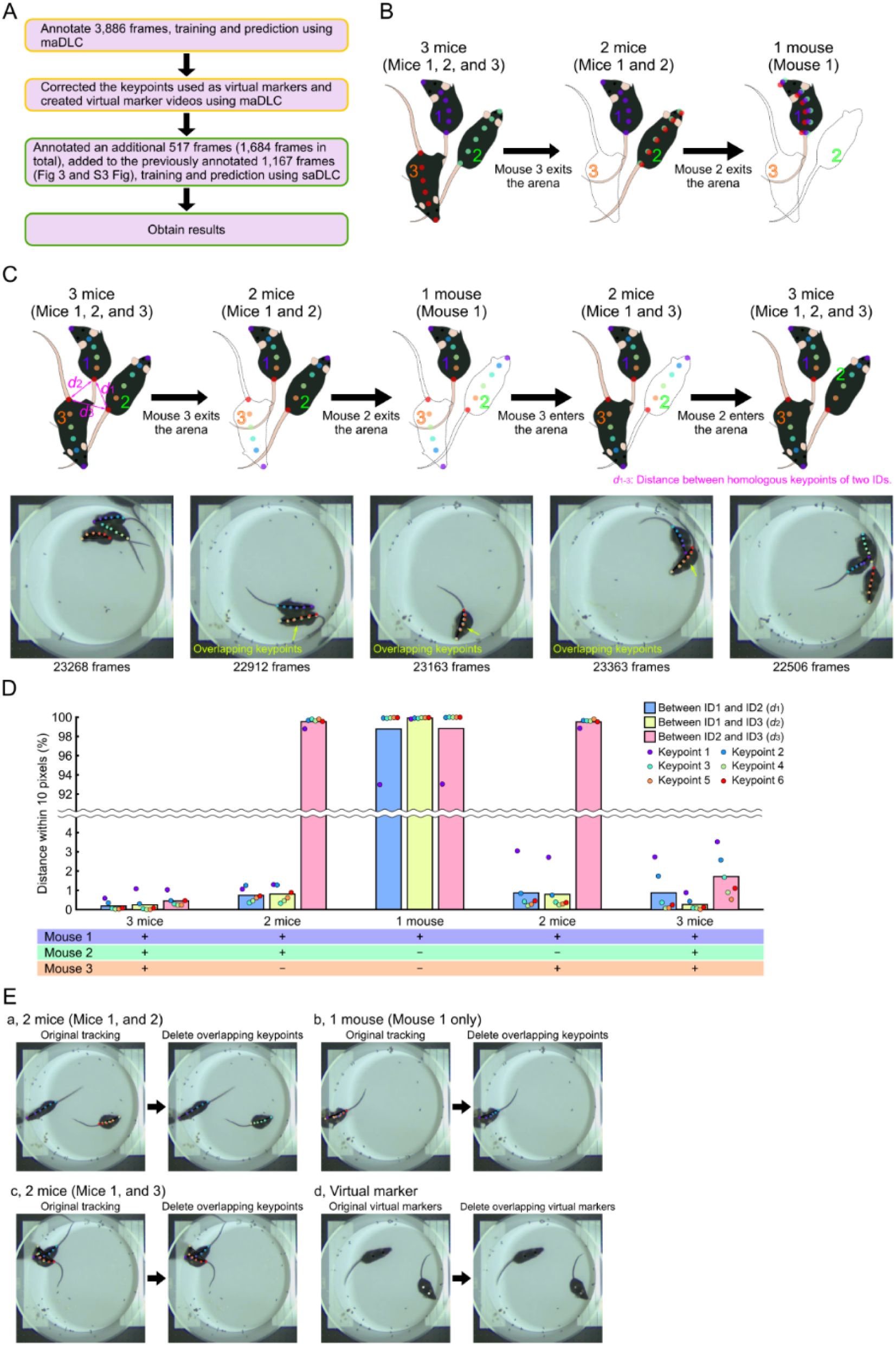
Tracking when the number of individuals in the video frame changes. (A) Overview of the verification process. The yellow box in the schematic represents the processes using multi-animal tracking tools, while the green box represents the processes using single-animal tracking tools. (B) Schematic diagram of the annotation method. The outlined mouse in the schematic indicates that the mouse was removed from the arena and is absent in the frame. In the case of three mice (left), six standard annotations were made for each mouse. In the case of two mice (center), keypoints for Mouse 2 and Mouse 3 were annotated on each body part of the same mouse. In the case of one mouse (right), keypoints for all mice were annotated on the body parts of that single mouse. Here, the labels are color-coded by individual. (C) Schematic diagram (top) of an experiment that altered the number of mice in the video frame by introducing or removing mice from the arena, and examples of tracking photos during this procedure (bottom). To verify tracking of other mice with leftover keypoints due to the absence of some mice, distances (d: pixels) between identical keypoints for three IDs were calculated for each keypoint. The stray keypoints indicated above the outlined mouse, which represents absence in the frame in the top schematic, were evaluated to determine whether they could predict another mouse in the arena, following the annotation method described in (B). Here, the labels are color-coded by body part. (D) If the distance between identical keypoints was within 10 pixels, the two ID keypoints were considered overlapping, and the frequency of such frames was calculated for each keypoint and ID pair combination. The diagram illustrates changes across different experimental conditions, with plots indicating the frequency for each keypoint and bars representing each ID pair. The “+” below the diagram indicates the presence of mice in the arena, while “−“ indicates their absence. Statistical analysis was conducted using the Friedman test for comparisons within each condition and the Kruskal–Wallis test for comparisons between conditions, with Bonferroni correction applied to account for the 45 total comparison pairs. Additionally, Cliff’s delta was calculated as an effect size for these pairs (S1 and S2 Table). (Ea-c) In Original tracking, markers for Mouse 1 (blue), Mouse 2 (green), and Mouse 3 (red) overlap under various conditions. After removing data from absent mice, a video with no overlaps and accurate tracking by correct IDs is created (Delete overlapping keypoints). (Ed) Even in markerless multi-animal pose tracking, markers can overlap. Removing such overlaps allows for creating virtual marker videos with accurate ID identification. In this example, extraneous white data obscuring the correct gray virtual marker are removed. The removal of unnecessary keypoint data is achieved through a Python code that allows specified ranges of tracking data obtained from DeepLabCut to be rewritten as NaN (no data) (S1 Protocol and S10C Fig).

**S6 Fig.**
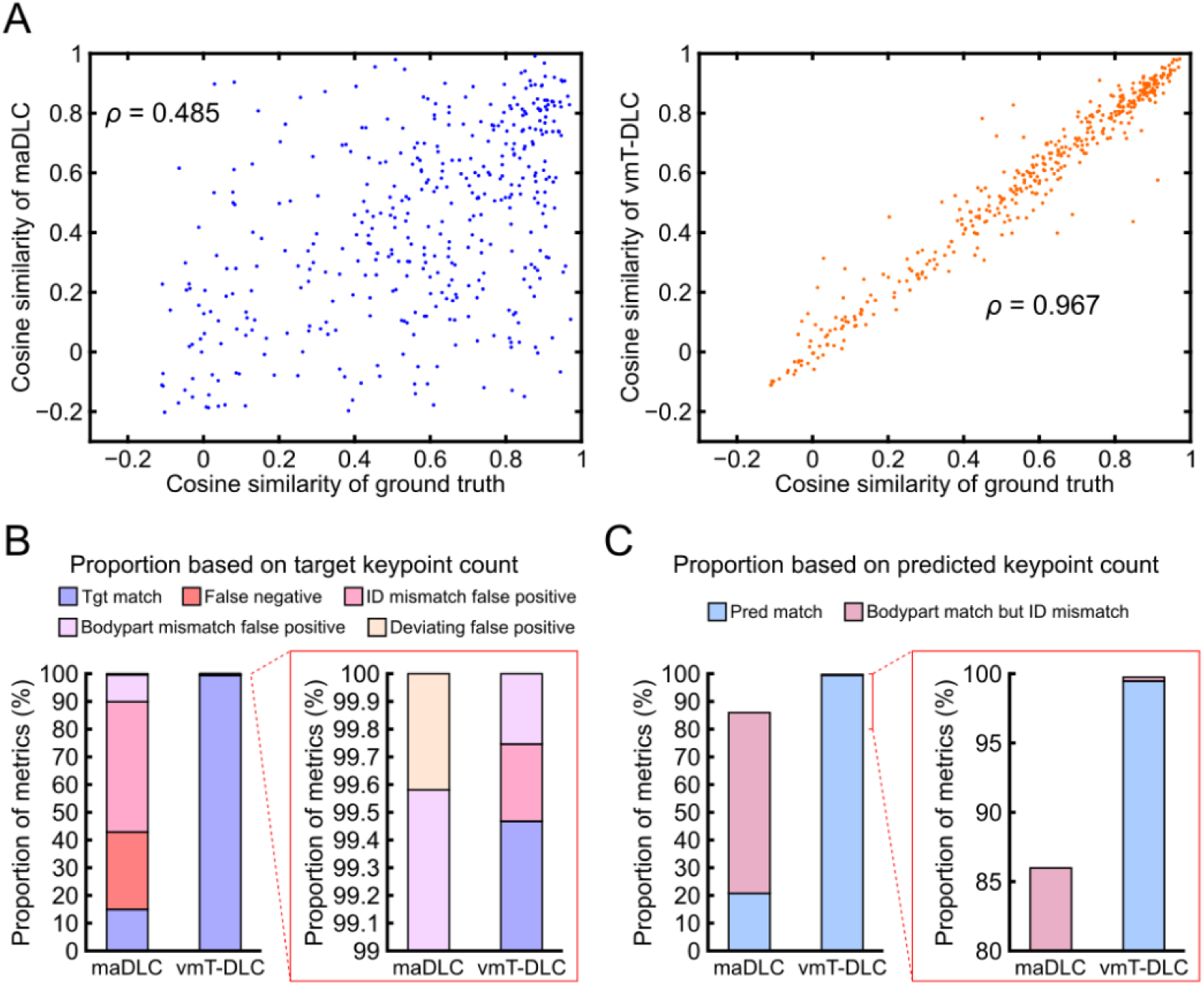
Tracking accuracy of 10 fish (Related to Fig 7). (A) Correlation of cosine similarity with the ground truth (GT). The GT consisted of 410 frames. The scatter plot on the left shows the cosine similarity between the ground truth and maDLC (*ρ* = 0.485, n =407), while the plot on the right shows the cosine similarity between the ground truth and vmT-DLC (*ρ* = 0.967, n = 407). Correlation analysis was conducted using Spearman’s rank correlation. (B) Stacked bar graph showing ground truth-based matches (Tgt match), false negatives, ID mismatch false positives (FPs), body part mismatch FPs, and deviating FPs for multi-animal DeepLabCut (maDLC) and virtual marker tracking with single-animal DeepLabCut (vmT-DLC). The graph within the red box is an enlarged view of the section with metrics showing low percentages. (C) Stacked bar graph of predicted keypoint matches (Pred match) and cases where body parts matched but the ID was mismatched (Bodypart match but ID mismatch). These values are derived from the data associated with the video evaluated in Fig 7.

**S7 Fig.**
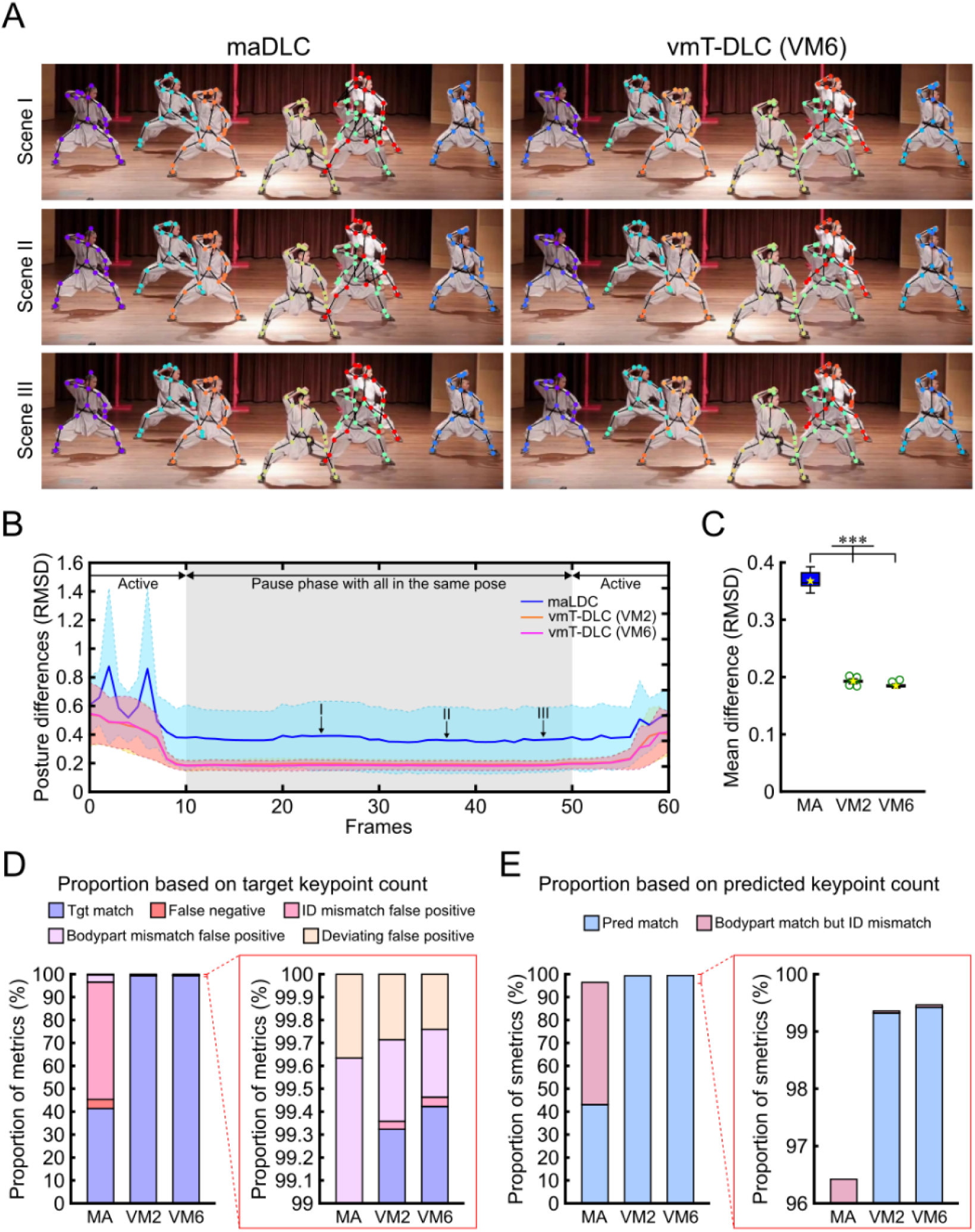
Tracking accuracy of multiple dancers (Related to Fig 8). (A) Tracking scenes using multi-animal DeepLabCut (maDLC) for markerless tracking and single-animal DeepLabCut for virtual marker tracking (vmT-DLC), with three examples where seven dancers are posed similarly and remain stationary for 2 s. The vmT-DLC results are based on the six-point virtual marker video (VM6). (B) Time series variation of RMSD over 3 s, including approximately 2 s of pause scenes. The shaded area along the plot line indicates standard deviation. The gray shaded area represents the pause scenes. I to III correspond to scenes I to III in (A). (C) Box plots of RMSD values for maDLC with markerless video (MA) and vmT-DLC with two-point virtual marker video (VM2) and VM6, calculated for all frames over a 2-s period, are presented. The bottom of the box represents the first quartile (Q1), the top represents the third quartile (Q3), the line inside the box indicates the median, the whiskers extend to 1.5 times the interquartile range (Q3-Q1), green circular markers denote outliers, and yellow star markers represent the mean. Statistical analysis was performed using the Friedman test with Bonferroni correction (n = 41). *: p < 0.05, **: p < 0.01, ***: p < 0.001. (D) Stacked bar graph showing ground truth (GT)-based matches (Tgt match), false negatives, ID mismatch false positives (FPs), body part mismatch FPs, and deviating FPs for MA, VM2, and VM6. The graph within the red box is an enlarged view of the section with metrics showing low percentages. (E) Stacked bar graph of predicted keypoint matches (Pred match) and cases where body parts matched but the ID was mismatched (Bodypart match but ID mismatch). The GT consisted of 150 frames.

**S8 Fig.**
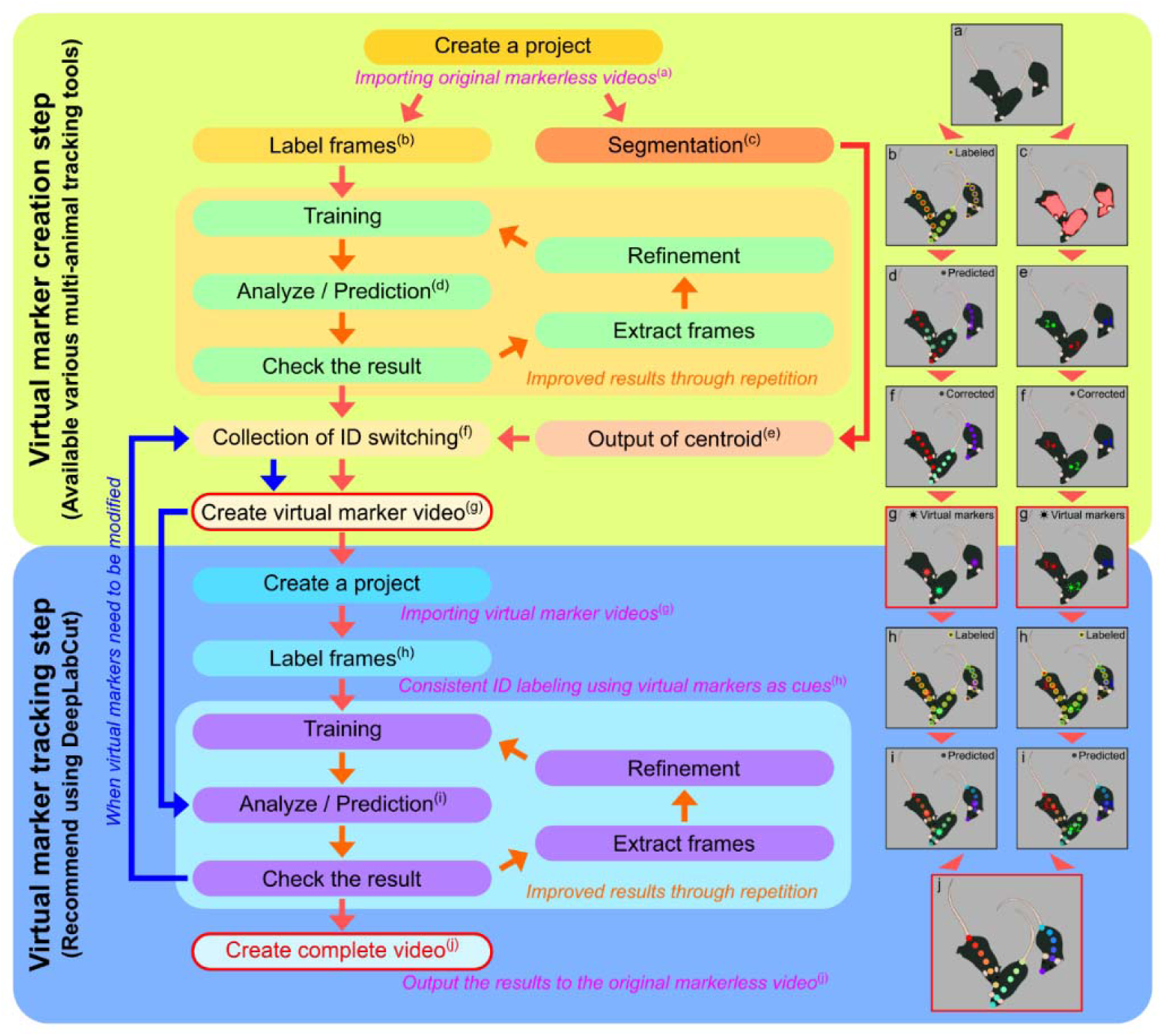
vmTracking Workflow. The vmTracking process consists of two major steps: creating the virtual marker video and tracking the virtual marker video. In the virtual marker creation step, a multi-animal tracking tool, such as multi-animal DeepLabCut (maDLC), Social LEAP (SLEAP), or idtracker.ai, is applied to the markerless video to track the animals. The tracking results are then corrected to ensure consistent ID assignment throughout the video, and the corrected tracking points are output as a new video. This video is called the virtual marker video. Thus, virtual markers are identification markers derived from the results of multi-animal tracking and do not physically exist. In the virtual marker tracking step, annotations are made so that consistent IDs are assigned to each individual across frames using the virtual markers as identification cues. Tracking is then performed using single-animal DeepLabCut, not maDLC. The resulting labels are applied to the original markerless video, producing a final tracking video without the virtual markers.

**S9 Fig.**
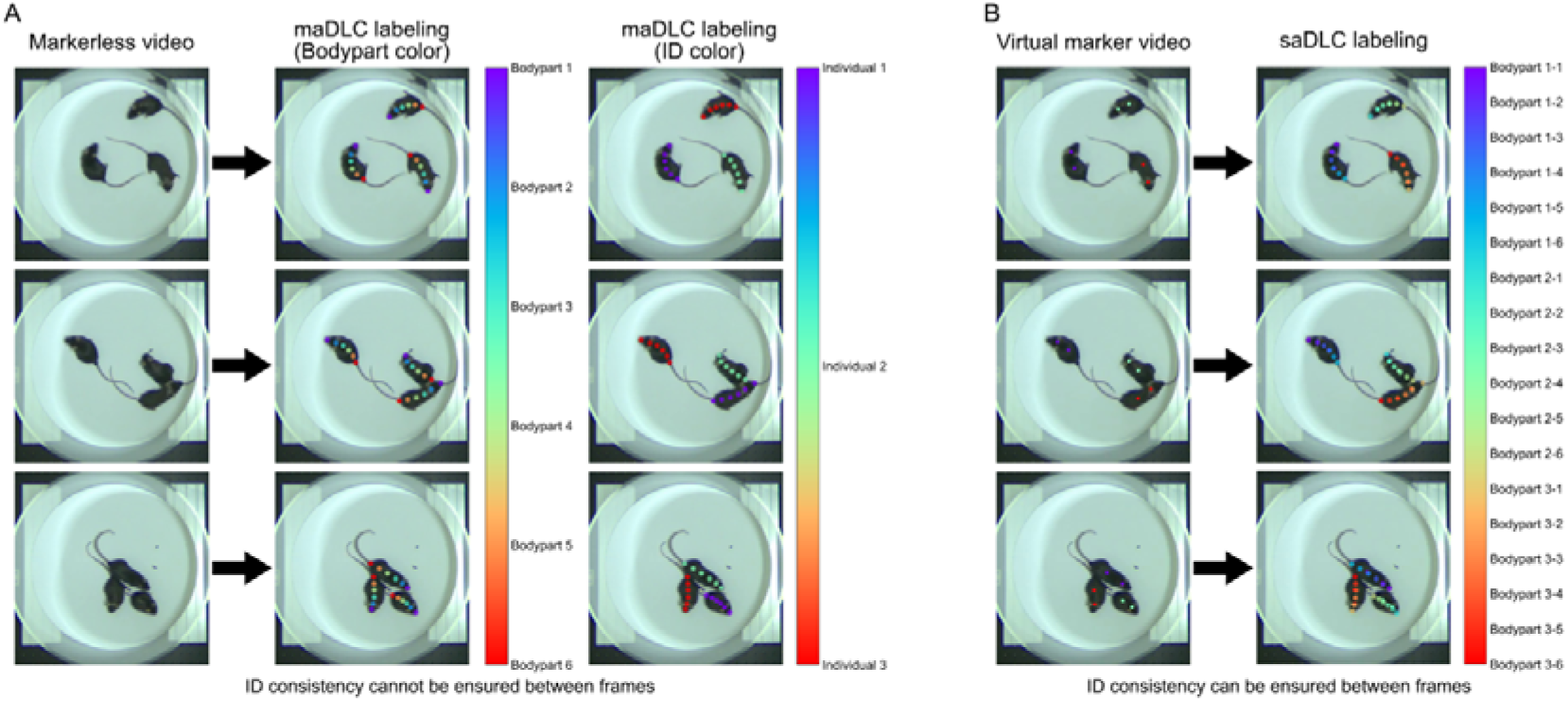
Comparison of annotations between markerless multi-animal tracking and virtual marker tracking (for DeepLabCut). (A) Example of annotations by multi-animal DeepLabCut (maDLC) for markerless videos. In maDLC, it is possible to display using either body part color or ID color modes, but in DeepLabCut 2.2, ID color does not appear to be displayed during the initial label frame step. In DeepLabCut 2.3’s napari-deeplabcut, the display mode can be switched using the “F” key. While it is essential to annotate all body parts for every individual, it does not matter which ID is assigned to which individual if the relationship between the individual and the ID cannot be determined in each frame. (B) Example of annotations by single-animal DeepLabCut (saDLC) for virtual marker videos. Since saDLC is designed for a single individual, the annotations assume that all body parts belong to one individual. The annotations are made so that consistent IDs are assigned to the same individual across frames, using the virtual markers as identification cues.

**S10 Fig.**
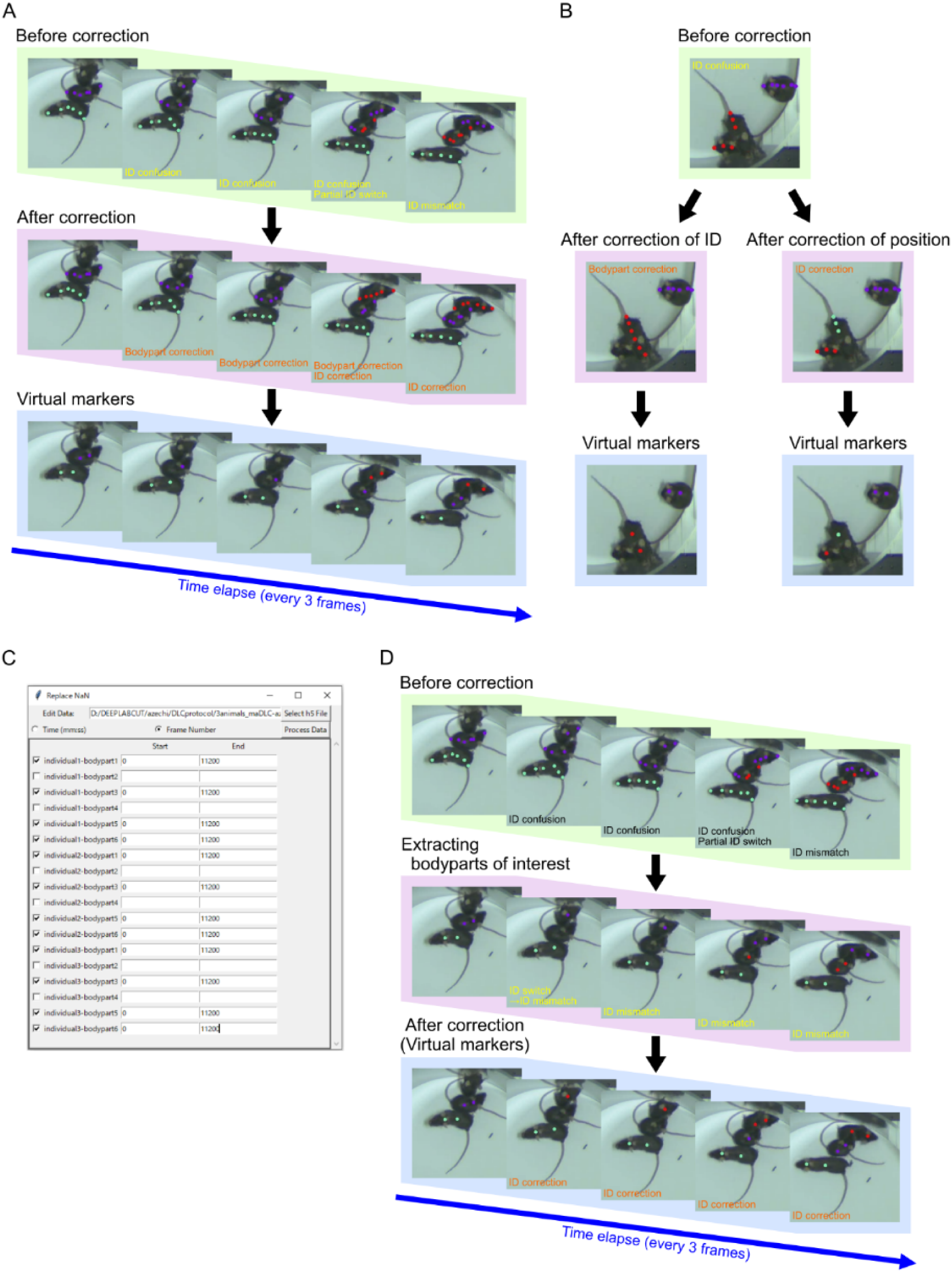
Correction of ID switches. (A) When ID switching (including partial switching) occurs, it is corrected. In this example, body parts that switched IDs (where purple labels were assigned across multiple individuals) were first adjusted so that all body parts were associated with a single individual. Then, the overall ID was corrected to ensure consistency throughout the video. These corrections can be made in DeepLabCut (DLC) using the “Refine tracklets” function (specific instructions available at: https://www.youtube.com/watch?v=bEuBKB7eqmk). (B) When a single ID label spans multiple individuals, two correction methods can be applied: adjusting body parts so that the label with the same color is assigned to one individual (left), or adjusting the ID to assign unique colors (IDs) to each individual (right). The choice of correction method affects the resulting virtual markers. While it is difficult to determine which approach is universally best, choosing a method that avoids biased ID omissions is recommended. (C) Screenshot of a Python-based GUI created to delete specified keypoints and data ranges from coordinate data files (in h5 format) obtained from DLC, replacing them with NaN (no data). Keypoints and data ranges for deletion can be specified by a combination of fps and time or by frame number. This tool enables users to retain only the keypoints designated as virtual markers and make necessary adjustments for virtual marker creation. (D) Corrections using the above-mentioned Python code when retaining only keypoints 2 and 4 in the same data as (A). In this example, the task is completed by correcting only ID switches, as non-virtual marker keypoints were removed. Clarifying which keypoints require correction is expected to improve the efficiency of the correction process.

**S11 Fig.**
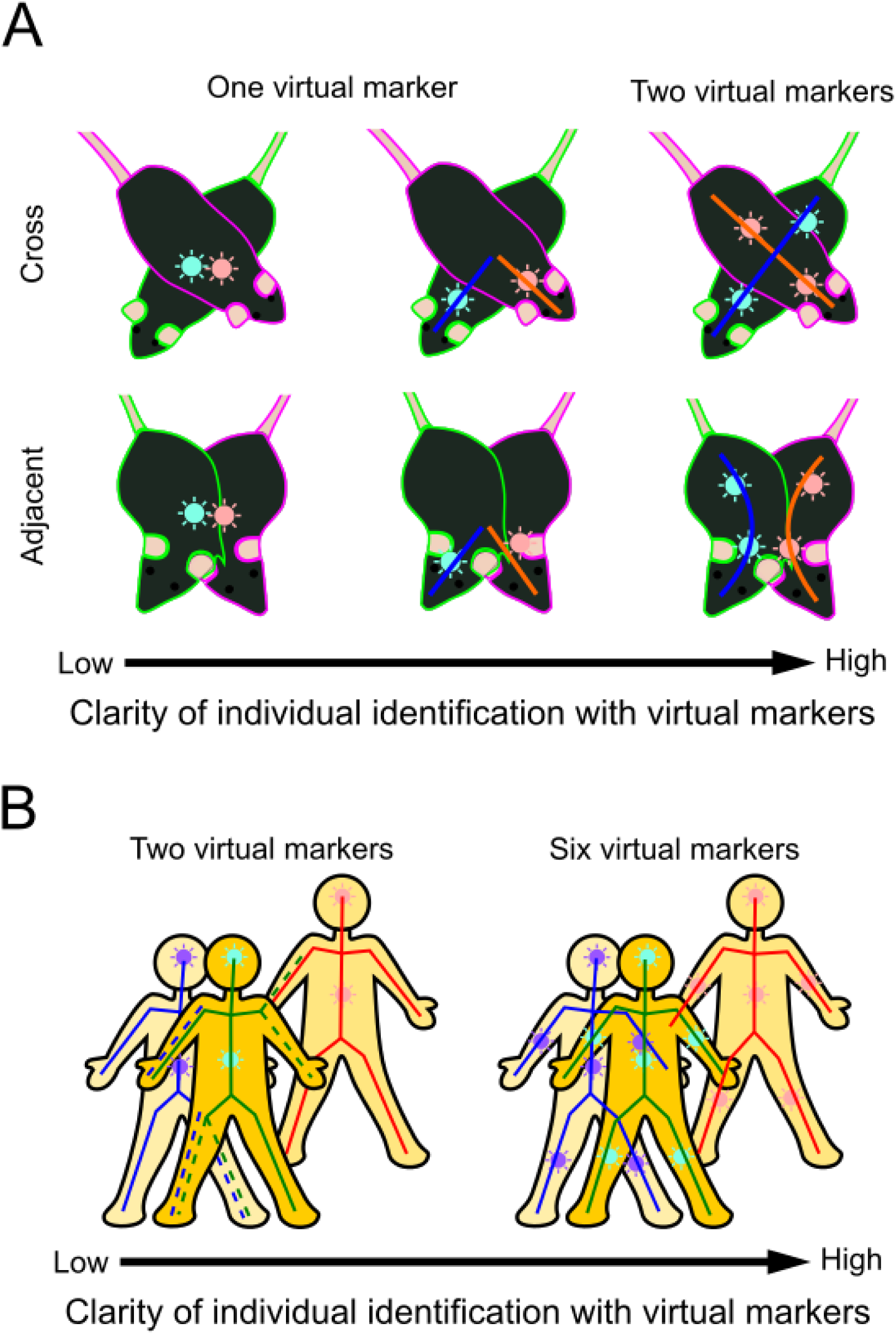
Considerations for the number and placement of virtual markers. (A) Comparison of cases where there is one virtual marker versus two virtual markers per mouse. The blue and orange solid lines represent the body axis that can be identified using the virtual markers. When one point is placed at the center of each mouse’s body, it becomes difficult to clearly distinguish between mice that are overlapping or adjacent (left). When one point is placed near the head, it becomes possible to identify individuals by their head position, but the area from the body to the tail cannot be clearly distinguished (middle). When two points are placed, one near the head and one near the tail, the two mice can be clearly distinguished (right). (B) Comparison of cases where there are two virtual markers versus six virtual markers per human. The blue, green, and red solid lines represent the predicted skeletons based on the virtual markers. When two virtual markers are placed on the head and abdomen of each individual, there is a high possibility that the arms and legs will be confused with those of other individuals (as indicated by the dotted lines). On the other hand, placing six virtual markers, including on the arms and legs, is expected to make it easier to detect the entire body, including the limbs, with the correct ID.

**S12 Fig.**
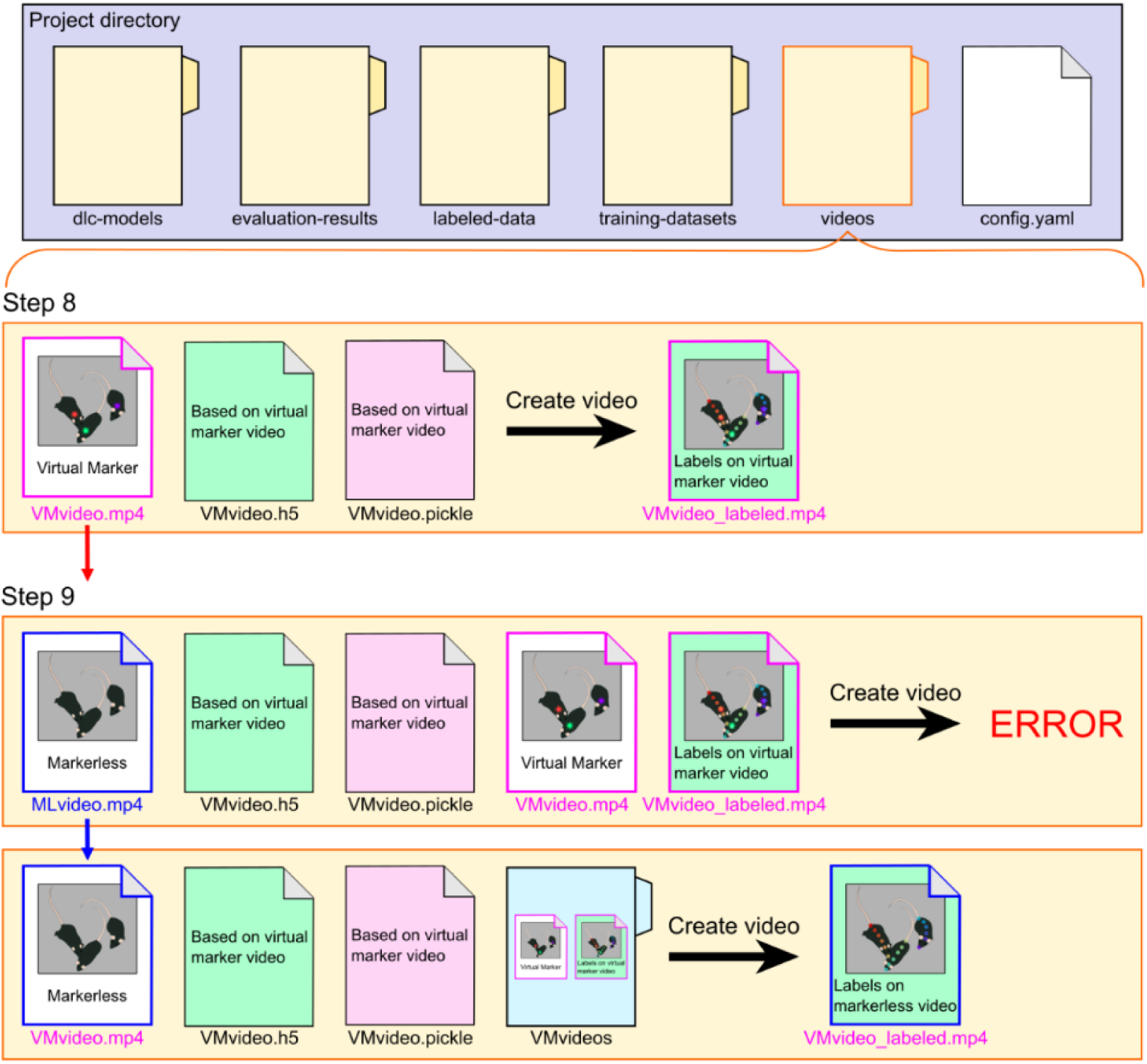
Outputting vmTracking results to the markerless original video. The top section of the figure is a schematic of the file structure within the project directory, followed by a schematic of the “videos” directory. When the results obtained from vmTracking are output to a video using the standard procedure, a video is created where the predicted labels are overlaid on the virtual marker video (Step 8). To apply the vmTracking results to a video without virtual markers, the results must be output to the markerless original video, and for this, the “videos” directory within the project directory needs to be properly prepared. If you simply place the markerless original video in the “videos” directory and attempt to create a video, an error will occur, and the video will not be generated (upper part of Step 9). If the videos used in Step 8 (the virtual marker video and the video with the overlaid results) are still in the “videos” directory, you should either rename these files or move them to a different directory (lower part of Step 9). Additionally, you need to rename the markerless original video appropriately. Since DeepLabCut processes files based on filename dependencies, the filename must correspond to the h5 and pickle files. Usually, changing the same filename as the virtual marker video used in vmTracking should work (lower part of Step 9). The Step numbers in the figure correspond to the steps in Protocol S1.

### Supplementary Table Legends

**S1 Table. Summary of statistical tests on the comparison of overlapping frame rates across different mouse count conditions for each mouse pair**

The lower-left values of the matrix represent the results of the Kruskal-Wallis test with Bonferroni correction, while the upper-right values represent Cliff’s effect size.

**S2 Table. Summary of statistical tests on the comparison of overlapping frame rates across different mouse pairs under various mouse count conditions**

The lower-left values of the matrix represent the results of the Friedman test with Bonferroni correction, while the upper-right values represent Cliff’s effect size.

**S1 Protocol. vmTracking protocol**

S1 Appendix. Example of bodyparts settings in the configuration file for multi-animal DeepLabCut

**S2 Appendix. Example of labeled data file in multi-animal DeepLabCut**

In an actual project, this file is created with the name CollectedData_experimenter.csv.

**S3 Appendix. Example of bodyparts settings in the configuration file for single-animal DeepLabCut for vmTracking**

**S4 Appendix. Example of labeled data file in single-animal DeepLabCut for vmTracking**

In an actual project, this file is created with the name CollectedData_experimenter.csv.

### Supplementary Video Legends

**S1 Video. Video clips demonstrating vmTracking applied across various species and conditions**

**S2 Video. Video clips demonstrating vmTracking applied to the tracking of five mice**

The comparative video clip demonstrated the performance of maDLC (A) and vmT-DLC (B) in tracking five mice. The video of vmT-DLC had virtual markers excluded. The video showed tracking results over 5 s at 0.1x speed. In the maDLC footage, there were identity switches indicated by label colors and missing predicted labels. In contrast, vmT-DLC consistently maintained accurate tracking throughout.

**S3 Video. Video clips demonstrating vmTracking applied to the tracking of three mice in an occlusion-crowded scene**

The comparative video clip demonstrated the performance of maDLC (A) and vmT-DLC (B) during the tracking of three mice in an occlusion-crowded scene. The video of vmT-DLC had virtual markers excluded. Beyond the 5-s mark in the video, the mouse labeled in red was partially obscured by the mouse labeled in green. In this scene, while the prediction of the mouse labeled in red was almost missing in maDLC, vmT-DLC maintained reasonable tracking.

**S4 Video. Video clips demonstrating vmTracking applied to the tracking of five mice in a non-occlusion-crowded scene**

The comparative video clip demonstrated the performance of maDLC (A) and vmT-DLC (B) during the tracking of three mice in a non-occlusion-crowded scene. The video of vmT-DLC had virtual markers excluded. In scenes without occlusion or crowding, both methods maintained consistent and accurate tracking.

**S5 Video. Video clips demonstrating the tracking of three mice with physical markers**

The comparative video clip demonstrated the performance of maDLC (A) and saDLC (B) during the tracking of three mice with physical marker. The video showed tracking results over 15 seconds at 0.5x speed.

**S6 Video. Video clips demonstrating vmTracking applied to the tracking of 14 lateralized keypoints in three mice**

The comparative video clip demonstrated the performance of maDLC (A), vmT-DLC (B), SLEAP top-down (SLEAP^TD^) (C), and vmT-LEAP (D) in tracking 14 lateralized keypoints in three mice. The video of vmT-DLC and vmT-LEAP had virtual markers excluded. The video showed tracking results over ten seconds at 0.25x speed.

**S7 Video. Video clips demonstrating vmTracking applied to the tracking of 10 fish**

The comparative video clip demonstrated the performance of maDLC (A) and vmT-DLC (B) in tracking a school of ten fish. The video of vmT-DLC had virtual markers excluded. The video showed tracking results over 15 s at 0.5x speed. In maDLC, there were occurrences of missing predicted labels and some keypoints predicting different fish, whereas vmT-DLC generally maintained stable tracking of each fish.

**S8 Video. Video clips demonstrating vmTracking applied to the tracking of 7 dancers**

The comparative video clip demonstrated the performance of maDLC (A) and vmT-DLC (B) in tracking seven dancers. The video of vmT-DLC had virtual markers excluded. In both maDLC and vmT-DLC, some keypoints occasionally predicted different dancers. However, while maDLC showed instances where entire sets of keypoints switched to tracking another dancer, vmT-DLC did not exhibit such switches.

