## Supplementary material for "vmTracking: Virtual Markers Overcome Occlusion and Crowding in Multi-Animal Pose Tracking": S1 Protocol

**Step-by-Step Protocol**

**Outline**

vmTracking is a method that achieves highly accurate markerless multi-animal pose estimation. In vmTracking, virtual markers are assigned to markerless animals to aid in individual identification and assist in multi-animal pose tracking. These virtual markers are labels generated from the results of existing markerless multi-animal tracking methods, which are then output onto markerless videos. Pose tracking is performed using a single-animal method on these videos with virtual markers, known as virtual marker videos. Therefore, vmTracking consists of two major steps: the generation of virtual marker videos (Virtual marker creation stage) and pose tracking using these videos (Virtual marker tracking stage). For multi-animal tracking to generate the virtual marker videos, tools such as multi-animal DeepLabCut (maDLC) or Social LEAP (SLEAP) can be used, as well as non-pose tracking tools like idtracker.ai. On the other hand, for the Virtual marker tracking stage, it is recommended to use single-animal DeepLabCut (saDLC). The entire workflow of vmTracking is shown in S8 Fig.

1. **Virtual marker creation step**
   1.1. **For DeepLabCut**

DeepLabCut can generally be executed by progressing through the GUI tabs from left to right. For details on installation and basic usage, please refer to the user guide (https://deeplabcut.github.io/DeepLabCut/README.html).

***Step 1: Create a multi-animal project (S8 Fig)***

Import the markerless video you want to track into DeepLabCut (S8a Fig) and create a multi-animal project. Important parameters to set in the configuration file include the number of animals to track (referred to as ‘individuals’ in the configuration file), the body parts you want to track (‘multianimalbodyparts’ in the configuration file), and the number of frames to extract for labeling (‘numframes2pick’ in the configuration file) (S1 Appendix). Other parameters should be adjusted according to the specifics of your experiment.

When setting up the body parts to track, it is acceptable to include only the body parts you wish to use as virtual markers. However, it is generally better to include as many body parts as possible. The number of extracted frames for labeling is optional, but it is expected that more frames will yield better results. Since the number of labeling frames can be added during the refinement process, we usually set it to 20–50 frames initially.

***Step 2: Label frames (Annotation, S8b Fig)***

After extracting the frames, proceed with labeling. While the method for frame extraction is optional, we usually use the automatic method via k-means clustering. During labeling, maintaining ID consistency across frames in markerless videos can be challenging. However, if identification is not possible, it is acceptable to label without maintaining ID consistency (S9A Fig). Once labeling is complete, a CollectedData file is created in the labeled-data folder within the project in both h5 and csv formats (S3 Appendix).

***Step 3: Training, evaluation, and analyzing videos (S8d Fig)***

Specify dlcrnet_ms5 as the network, create the training dataset, and begin training. We set the maximum number of iterations for training to 200,000, which is the upper limit for multi-animal DLC. Under this condition, training takes several hours. Once training is complete, proceed to evaluation and then to video analysis. After obtaining prediction results from video analysis, merge these results with the original video to create a labeled video and visually inspect the predictions.

If there are frequent ID switches, we recommend extracting outlier frames (we typically use the jump method) for refinement, followed by re-training. Don't forget to merge the dataset after labeling. When the dataset is merged, the number of iterations (different from the maximum number of iterations set for training) in the configuration file is updated, allowing you to re-train the model with the dataset that includes the newly added labeling frames. If you choose not to re-train, proceed to the next step.

***Step4: Correction of ID switching (S8f Fig)***

maDLC stitches together tracklets, which are fragments of motion trajectories obtained from video analysis, to form continuous behavioral trajectories for each animal (7). However, the generation of tracklets can be interrupted by occlusion, leading to failures in stitching and resulting in ID switches. Manually correct the obtained prediction results. maDLC includes a ‘Refine Tracklets’ function, which allows you to correct ID switches and manually adjust keypoints. In short, to correct ID switches, place flags at the start and end points of the range you want to adjust, then press the Lasso button. Use it to select the keypoints you want to swap, and choose the desired color from the list next to the individual name at the top of the screen. To move individual keypoints, press the Drug button, which enables dragging individual keypoints (for details on the correction method refer to: https://www.youtube.com/watch?v=YSRQT8N2vFE&t=11s). Focus on correcting ID switches for the keypoints intended to be used as virtual markers, ensuring they are consistently tracked with the same ID throughout the video (S10 Fig). If the predicted points are detected on the correct individual, slight misplacements (e.g., a keypoint on the torso being predicted as the head) are not a significant issue. Important corrections include fixing ID switches that occur during a sequence of movements (S10A Fig) and resolving cases where predicted labels are spread across multiple individuals (S10B Fig). In the latter case, either correct the ID for some predicted points or properly assign the predicted labels to a single individual. However, there is no single correct approach. Additionally, it should be noted that in maDLC, predictions may sometimes be missing, and certain individuals may be completely overlooked in specific frames. The Refine Tracklets feature cannot compensate for these missing predictions.

In this step, it is actually sufficient to correct only the keypoints used as virtual markers. However, in reality, there are keypoints that are not intended to be used as virtual markers, and therefore, they can obstruct the correction process. To address this, you can use our Python code (S10C Fig, and available on Zenodo) to exclude data related to unnecessary keypoints, making it easier to focus on the keypoints intended for use as virtual markers. This code reads an h5 format file of coordinate data and allows you to replace the specified range of data with NaN by specifying either the range of video time or the range of frame rate and frame number. The selected file is designed to be backed up before any modifications are made.

Although a virtual marker can consist of just one keypoint, since maDLC predictions can sometimes be missing, using multiple points as virtual markers helps avoid cases where no virtual marker is applied. For example, if two points are set as virtual markers and one is missing, the other can still function as a virtual marker. Additionally, using two markers allows for better determination of the body axis, which is especially useful when two individuals overlap, enabling more accurate identification (S11A Fig). This also reduces the chance of misidentification during labeling by the experimenter. In cases such as humans, where arms or legs may be in close proximity and their IDs could partially switch, placing virtual markers on the arms or legs can help achieve more accurate identification of the entire individual (S11B Fig).

Our tests have shown that even with fewer virtual markers, the performance is significantly better than traditional markerless multi-animal pose tracking (Fig 5, 8, and S7 Fig), and considering the effort required for corrections, increasing the number of virtual markers may not be very meaningful. However, as mentioned above, using multiple virtual markers can indeed improve tracking accuracy to some extent (Fig 5, 8, and S7 Fig). While the overall benefits may be minimal, adding markers could still be effective in situations where ID switching is more likely to occur. On the other hand, it may be advisable not to add too many virtual markers. We consider virtual markers primarily as clues for individual identification, and as long as they serve that purpose, it is better for the virtual markers to be moderately ambiguous, even allowing for some marker loss (Fig. 2I, 3I, S3I). Therefore, we take care to ensure that the physical characteristics of the target are not obscured by the virtual markers.

For example, when there are too many virtual markers, they may become indicators of specific body parts rather than simply being identifiers for individual recognition. As a result, the dependence on virtual markers for tracking may become excessive. The more consistent the location of the virtual markers is, the more accurately the relationship between the virtual markers and the predicted positions will be learned, potentially leading to such issues. This could also be something to be cautious of even with fewer markers. To some extent, virtual markers inevitably serve as clues to body parts. For instance, if a keypoint on the head is used as a virtual marker, it naturally results in the virtual markers being predominantly located around the head. In such cases, if a virtual marker happens to be placed far from the head, it may lead to an incorrect prediction of that position as the head. It is believed that having a less rigid location for virtual markers can reduce such occurrences. Therefore, making fine corrections to the position of body parts during the virtual marker creation step may have little advantage in terms of both the effort involved and the accuracy of vmTracking.

***Step 5: Create a virtual marker video (S8g Fig)***

Using the corrected data, create a labeled video (virtual marker video) that outputs only the keypoints intended for use as virtual markers. When creating the video, choose the output method that uses ID colors, so that the IDs are color-coded in the video. The size of the virtual markers does not need to be excessively large; it just needs to be large enough to be visible.

1.2. **For SLEAP**

While the basic workflow for using SLEAP is similar to DeepLabCut, there are some operational differences that will be briefly explained. For details on installation and basic usage, please refer to the user guide (https://sleap.ai/).

***Step 1: Create a project for multi-animal tracking (S8a Fig)***

From the Videos tab on the right side of the GUI, add the markerless video you want to analyze. In the Skeleton tab, set the Nodes (equivalent to ‘bodyparts’ in DeepLabCut) and, if necessary, the Edges (equivalent to ‘skeleton’ in DeepLabCut). It is acceptable to set Nodes only for the body parts you intend to use as virtual markers.

***Step 2: Label frames (Annotation, S8b Fig)***

After extracting frames from the Labeling Suggestions tab, proceed with labeling. The method of frame extraction is optional, but extracting around 20–50 frames using the sample method should suffice. During labeling, maintaining ID consistency across frames in markerless videos can be challenging. However, if identification is not possible, it is acceptable to label without maintaining ID consistency. You can also import and use annotation data from DeepLabCut.

***Step 3: Training and inference (Equivalent to DeepLabCut's training, evaluation, and* aanalyzing videos, S8d Fig)**

Training and prediction follow the basic analysis procedures described in the user guide. There are two pipelines for multi-animal analysis: top-down and bottom-up. Among the parameters that need to be set, specify the number of Max instances if known, and set the other parameters according to your experiment. SLEAP’s training and prediction process is typically faster than DeepLabCut. Once training is completed, proceed with inference to obtain prediction results. After obtaining the prediction results for the entire video, use the seek bar at the bottom of the GUI to visually inspect the predictions. If frequent ID switches occur, extract the frames with poor predictions, correct them, and re-train the model.

***Step 4: Correction of ID switching (S8f Fig)***

Manually correct the obtained prediction results. As with DeepLabCut, focus on correcting ID switches for the keypoints intended to be used as virtual markers, ensuring consistent ID tracking throughout the video. In SLEAP, you can easily correct ID switches by selecting the instance you want to swap and using a shortcut key operation (Ctrl + number key). Keypoints can be moved by selecting the keypoint you wish to move and dragging it. Additionally, SLEAP allows you to supplement missing nodes in the predictions. Moreover, you can remove unnecessary nodes in the Skeleton tab of the GUI, leaving only the nodes intended for use as virtual markers.

***Step 5: Create a virtual marker video (S8g Fig)***

Using the corrected data, create a labeled video (virtual marker video) that outputs only the keypoints intended for use as virtual markers. Ensure that the IDs are color-coded by selecting ‘Apply Distinct Colors To’ and choosing ‘instances’ from the view menu before creating the video. Additionally, do not output the edges.

1.3. **For other multi-animal tracking tools**

In addition to the multi-animal pose tracking tools mentioned above, we have confirmed that idtracker.ai can also be used for creating virtual markers. idtracker.ai tracks the center of mass of individuals identified through segmentation (S8c Fig), and after correcting any ID switches (S8e Fig), this centroid (along with the ID number) serves as the virtual marker. Unlike DeepLabCut or SLEAP, there are no special techniques required for creating virtual markers with idtracker.ai; simply applying the standard method as described in the user guide (https://idtracker.ai/latest/) is sufficient, so further explanation is omitted here. Although we have not tested them, it is likely that other similar multi-animal tracking tools can also be used to create virtual markers.

2. **Virtual marker tracking step**

This step will only explain the method using saDLC, which, based on our verification results, can achieve high accuracy across all videos.

***Step 6: Create a single-animal project***

Create a single-animal project to analyze the virtual marker video created in the previous step. In the configuration file, set the number of keypoints to track as the number of animals multiplied by the number of body parts to track, and name them accordingly (e.g., 1-1, 1-2, 1-3, 2-1, 2-2, 2-3, etc., S2 Appendix). The number of extracted frames is optional, but more frames typically yield better results. We usually set it to 20–50 frames. Additionally, set the skeleton if necessary. Other parameters should be adjusted according to the specifics of your experiment.

***Step 7: Label frames (S8h Fig)***

After extracting the frames, label them. We typically use the automatic method via k-means clustering for frame extraction. During labeling, use the virtual markers attached to the image as cues for individual identification, ensuring ID consistency across frames (e.g., consistently labeling the individual with a purple virtual marker as ID1, S11B Fig). Once labeling is complete, a CollectedData file is created in the labeled-data folder within the project in both h5 and csv formats (S4 Appendix).

***Step 8: Training, evaluation, and analyzing videos (S8i Fig)***

Create the training dataset and start training. The choice of network for creating the training dataset is flexible, but we use EfficientNet (specifically, EfficientNet-b0 as implemented in version 2.2.3 for this study). While a higher number of iterations may yield better results, we usually set the maximum number of iterations to between 200,000 and 500,000, which should be sufficient. Once training is complete, proceed to evaluation and then video analysis. After obtaining the prediction results from video analysis, merge these results with the virtual marker video to create a labeled video and visually check the predictions. Since it is unlikely to achieve satisfactory accuracy with the initial training, we recommend extracting outlier frames (we usually use the jump method) for refinement, followed by re-training. During this refinement, adjust the labels using the virtual markers as a reference. Don’t forget to merge the dataset after refinement.

If you notice any virtual marker ID switching or other necessary corrections during the video check, return to step 4 of the virtual marker creation step, correct the relevant parts, recreate the virtual marker video, and reanalyze the video.

***Step 9: Create a complete video excluding virtual markers (S8j Fig)***

Once you have repeated the training and achieved satisfactory results, you can obtain a tracking video without virtual markers by merging these results with the original video. Since DeepLabCut’s labeled video creation process relies on file naming dependencies, it is important to prepare the videos directory within the project appropriately before proceeding. Specifically, place the original markerless video in the videos directory and rename it to match the filename of the virtual marker video. In doing so, you will need to either rename the previously existing virtual marker video or temporarily move it to another directory (S12 Fig).
